# Flux-centric coarse-graining of hepatocyte metabolic network captures acute-on-chronic liver failure dynamics in alcohol-associated hepatitis

**DOI:** 10.64898/2026.11.26.769501

**Authors:** Madison Shoraka, Rajanikanth Vadigepalli

## Abstract

Acute-on-chronic diseases are marked by progressive deterioration bifurcated by sudden shifts in the compensatory state. Alcohol-associated hepatitis (AH) is an acute manifestation in patients with chronic alcohol-associated liver disease. Metabolic decompensation is a hallmark of AH. How the dysfunction of multiple metabolic pathways collectively lead to overall decompensation remains poorly understood. We developed a flux-centric, coarse-grained mathematical framework to represent hepatocyte metabolism as a network of metabolic modules operating under a constrained resource. We define the module-level metabolic flux as the overall rate at which metabolites are processed through the corresponding reactions. The stoichiometry of shared resources determines how the module-level fluxes re-balance upon perturbation to the network. We included four interacting modules: glutathione synthesis, methionine cycle, urea cycle, and fatty acid biosynthesis, which are coupled through shared metabolites. We simulated a range of liver states under varying ethanol intake episodes corresponding to healthy and disease conditions. We examined the impact of corticosteroid treatment on the metabolic network in severe AH cases. Simulation results were consistent with clinical observations including down-regulation of ammonia detoxification, reduced antioxidant capacity due to depletion of mitochondrial glutathione, up-regulated fatty acid biosynthesis, and decreased methylation potential. Relaxing overall resource constraints by modifying the network coupling significantly reduced the consistency with clinical phenotypes. The emergent dynamics of the metabolic network captured the heterogeneity of response to repeated ethanol intake and corticosteroid treatment. Simulated corticosteroid treatment scenarios yielded a 72-82% response rate for severe AH cases consistent with the average clinical response rate of 75%. The simulations also suggest that the treatment can destabilize severe AH cases with milder metabolic decompensation. Overall, our model-based analysis highlighted how decompensation can emerge from interacting metabolic cycles under resource constraints in AH. This mathematical framework is broadly applicable to characterize the landscape of metabolic decompensation in acute-on-chronic failure scenarios.

**Author summary:** Alcohol-associated liver disease (ALD) develops quietly over years of drinking, often without symptoms, until the liver suddenly fails. This crisis, alcohol-associated hepatitis (AH), carries a 30-50% risk of death in the short term, and the standard treatment, corticosteroids, helps only about three in four patients. Doctors have few ways to tell who is about to tip into failure or who will respond to treatment. Answering these questions means understanding how the liver’s many biochemical processes interact under the strain of alcohol intake, but detailed kinetic models of metabolism need measurements that cannot be obtained from patients. We instead built a simplified model of four liver metabolic processes (antioxidant production, clearing ammonia, methyl donor production, and making fat) that compete for a shared, limited pool of resources. Like springs tied together, pulling on one shifts all the others. The model reproduces what is seen in patients: rising ammonia, fat buildup, sudden rather than gradual failure after repeated drinking, and the roughly 75% corticosteroid treatment response rate. The same mathematical approach could be applied to other diseases marked by a long, silent buildup of dysfunction and a sudden crisis of failure.

## 1 Introduction

Alcohol-associated liver disease (ALD) is the most prevalent form of chronic liver disease worldwide, developing over years of sustained alcohol exposure and advancing through a well characterized histological spectrum from alcohol-associated fatty liver (steatosis) to steatohepatitis, and subsequently to fibrosis, cirrhosis, and, in some cases, hepatocellular carcinoma [1]. Throughout much of this progression the disease remains clinically undiagnosed, and patients frequently come to attention only upon acute decompensation. Its most severe manifestation, alcohol-associated hepatitis (AH), carries short-term mortality of approximately 30-50% [2] and is defined clinically by an intense hepatic and systemic inflammatory response superimposed on chronic injury, which precipitates rapid hepatic decompensation [3, 4]. The underlying pathogenesis is multifactorial, including hepatic steatosis, oxidative stress, acetaldehyde mediated cellular toxicity, and cytokine and chemokine induced inflammation [1]. Despite broad recognition of these processes, once AH occurs there are few effective therapeutic options and limited ability to predict which patients will progress to decompensation [1, 3–6]. The current standard of care for AH is the corticosteroid treatment, prednisolone. The response to corticosteroid treatment is heterogeneous, with roughly 75% of eligible patients demonstrating a favorable short-term response and the remainder showing no benefit or even deterioration [7].

Metabolic reprogramming is a hallmark of functional decompensation in AH [1]. During the progression of ALD, glucose use is remodeled away from maintaining glycogen homeostasis [8]. Hepatic glutathione (GSH), the primary intracellular antioxidant and especially concentrated in hepatocytes, is broadly dysregulated and depleted [9, 10]. Methionine metabolism is likewise disrupted: ethanol and its reactive oxygen species (ROS) products inhibit key enzymes of the methionine cycle whose loss simultaneously impairs antioxidant defense and genetic regulation [11–13]. Urea cycle function is similarly compromised, with hyperammonemia as the downstream consequence [14, 15]. The impairment of these metabolic processes crosses a critical thresholds for maintaining liver metabolic function [16]. In addition to the above metabolic modules’ dysregulation, ethanol also promotes fatty acid biosynthesis and impairs *β* oxidation, making hepatic steatosis a near universal early feature in ALD [17–19]. Such studies have led to multiple AH treatments being explored with varying degrees of success including zinc [20] and GSH precursor supplementation [21]. There remains a gap in identifying effective therapeutic targets that can rescue metabolic decompensation.

Systems level and multiomics approaches have been applied to study hepatic metabolism as it relates to AH [8, 22, 23]. Such studies build on genome scale human metabolism reconstructions: Human1, a pioneering application, enumerates the entire human cell metabolism down to individual biochemical equations and their associated gene regulators [24]. Gille and colleagues developed HepatoNet1, a comprehensive reconstruction of the human hepatocyte encompassing over 2500 reactions [25].

Complementing these, multi-layer regulatory models integrate control across levels of biological organization, as in SteatoNet, the first multi-layered model of hepatic metabolism to incorporate transcriptional, enzymatic, and metabolic regulation, used to examine the integrated effects of lipid metabolic perturbations on liver pathology [26]. These frameworks share a common insight: single pathway analysis cannot capture the emergent behavior of a system in which dozens of metabolic cycles interact through shared metabolites. A recent study extended this insight to an organismal scale, reviewing how ethanol induced cascades propagate through interconnected metabolic and inflammatory networks to disrupt homeostasis across liver, heart, pancreas, and brain [16]. Our group has applied this genome scale metabolic modeling and analysis approach to AH. Through integrated metabolomic and transcriptomic analysis of AH liver tissue we uncovered a profound reprogramming of hepatic glucose metabolism [8]. Our studies additionally revealed sequential dysregulation of glutathione metabolism as one of the earliest detectable signatures of AH progression [22]. In our recent *in silico* study, we identified multi-hit genetic perturbations that are likely to reset the glycolytic component of patient specific metabolic reprogramming in AH [23].

In the present study, we explore how dynamic interdependencies between metabolic modules drive the metabolic decompensation characteristic of AH. Human metabolism is a high dimensional nonlinear system with compartmental transport and multi-layered regulation, and its kinetic scale simulation is limited by parameter non-identifiability, numerical stiffness from widely separated reaction timescales, and sensitivity amplification from strong interdependencies. These limits are especially acute for AH, where the required data are not readily available: there is no faithful animal model, and the enzyme resolved time course measurements a kinetic model would demand are not collected from patients. We therefore develop a hybrid model, combining flux based analysis from genome-scale models and the governing principles from kinetic models. We analyze four interacting metabolic modules: glutathione synthesis, the methionine cycle, the urea cycle, and fatty acid biosynthesis. Each module is independently implicated in AH and is a dominant flux handle on the metabolites produced or affected by ethanol. To govern how these fluxes redistribute, we draw on metabolic control analysis, which formalizes the principle that flux changes across a network are relatively constrained rather than independent. In our scheme, this constraint takes the form of a shared pool of metabolic control capacity: as each module is perturbed over time, the pool dictates how the modules adapt and reallocate flux in response.

We use this modeling and analysis approach to assess (1) whether coarse-grained flux tradeoffs reproduce known AH phenotypes as an emergent consequence of metabolic resource competition under an overall adaptation constraint, (2) whether repeated acute-on-chronic ethanol exposure produces a qualitative transition to decompensation rather than a continuous decline, and (3) whether network linkage and metabolic flux constraints can explain the heterogeneous response to standard of care corticosteroid treatment to AH.

## 2 Results

We developed a mathematical formalism for metabolic cycles which we applied to study hepatic metabolism whose effective velocities, rather than metabolite concentrations, are the primary dynamical variables. The overarching principle is drawn from metabolic control analysis (MCA), which holds that control over flux is distributed across a pathway and constrained rather than independent. In MCA, this appears as the summation theorem: flux control coefficients sum to one, so gaining control at one step means losing it elsewhere. Our framework makes this constraint explicit as a fixed total velocity shared across metabolic modules. Perturbation cannot create capacity; it can only reallocate it. Each tracked metabolite is linked to the modules that produce or consume it through a stoichiometric coupling matrix. From the steady state we derive capacity fractions, the share of each metabolite’s baseline turnover handled by each metabolic module. Like control coefficients in MCA, these capacity fractions in our formalism add up to one across metabolic modules. Metabolite variables act as load signals, in the spirit of MCA elasticities: each quantifies how a given deviation from baseline in metabolite concentration exerts demand on its associated metabolic cycles. The unmet demand of metabolic modules forces the reallocation of shared reactants and cofactors, such as glutamate, acetyl-CoA, and NAD^+^*/*NADH as defined by the module adjacency network (Fig. 1).

**Fig 1.**
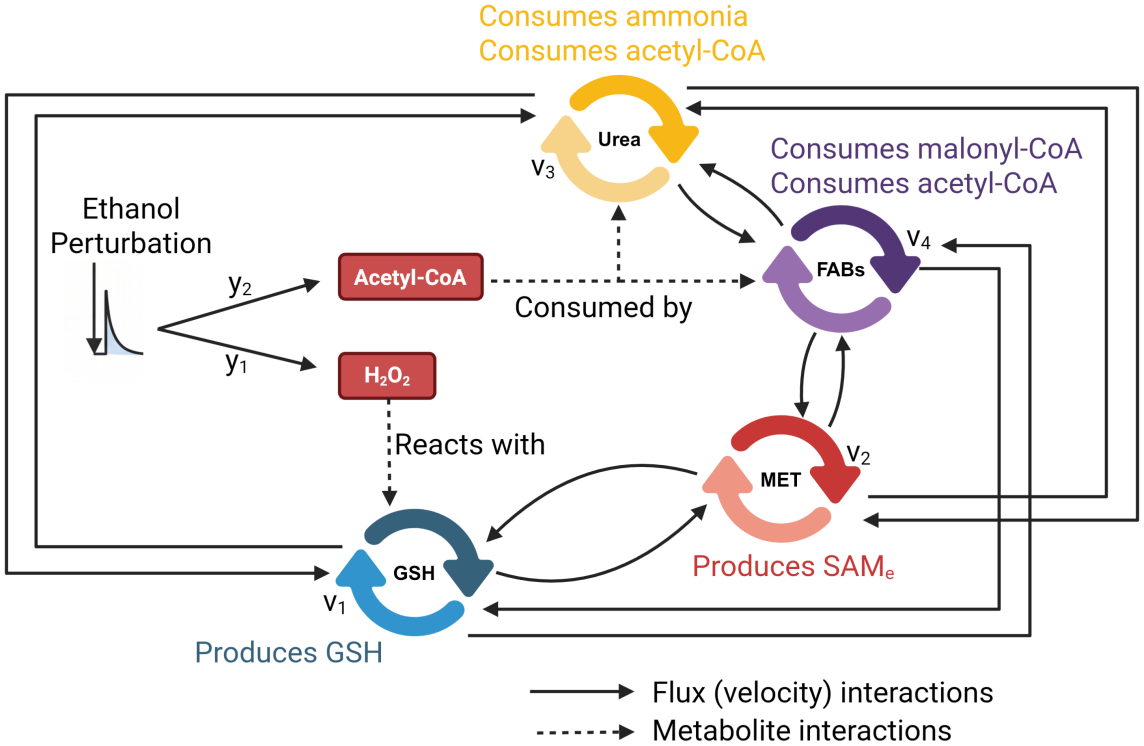
Specifications for ethanol perturbations: Ethanol propagates through our system, yielding H_2_O_2_ (*y*_1_) via oxidation and acetyl-CoA (*y*_2_) further downstream. The metabolic cycles are connected via an adjacency matrix as defined by their composing biochemical reactions. The perturbations are then permitted to flow through the system as it attempts to re-stabilize. We define a failed system as that which under perturbation has a transient spike in H_2_O_2_ concentration that is not metabolized by GSH and a metabolically decompensated system as that whose fluxes are unable to re-stabilize within a given time frame.

We applied this formalism to elucidate the underlying mechanisms of alcohol induced hepatitis (AH) under the progression of alcohol-associated liver disease (ALD). To capture these dynamics, we chose to model four metabolic modules: the glutathione synthesis cycle, the methionine synthesis cycle, the urea cycle, and even-chain fatty acid biosynthesis (FABs). See Sec. 3.1 for further details on how these cycles capture AH and are modified under ALD progression. Ethanol then enters the dynamics as an exogenous, zero order cleared input that generates mitochondrial H_2_O_2_, which drains glutathione through fast third order kinetics, and acetate, which feeds acetyl-CoA. Two patient specific yield coefficients and a transport coefficient set the strength of this forcing. Because capacity is conserved and must be reallocated through a coupled network, pathway level phenotypes such as suppressed urea flux and enhanced lipogenesis arise from competition among modules rather than being imposed directly. Oxidative and metabolic failure appear as threshold losses of stability in the coupled dynamics.

The central hypothesis we seek to test is that the emergent phenotypes of AH arise from the interlinkage of these subsystems and their competition for shared resources, rather than from the additive failure of individual pathways. We therefore organize the results as a series of concordance comparisons, each pairing a clinical or experimental scenario with the corresponding model behavior. Because longitudinal intrahepatic metabolomics at the hours timescale does not exist for AH patients, validation is conducted not against fine-grained temporal trajectories but at the level of emergent behavior: steady-state shifts, failure frequencies, and treatment response rates that can be compared against clinical and experimental observation. This is consistent with validation practice in other reduced order biological models operating beyond the reach of current measurement technology. Without hard coding any pathway specific failure, we show that this construction reproduces the canonical behaviors of ethanol stressed hepatic metabolism: episodic oxidative stress following alcohol intake, delayed mitochondrial acetyl-CoA accumulation, competition between detoxification pathways under limited capacity, and progressive depletion of mitochondrial glutathione.

### 2.1 Concordance with the canonical clinical phenotypes

#### Clinical scenario

A range of patient profiles, spanning the physiological parameter space above, each receive an acute ethanol dose. Clinically, ethanol metabolism is characterized by four key attributes:

1. **Decreased methylation potential**. In the model, this would be indicated by a decrease in the total velocity of the methionine cycle over time. [13]
2. **Decreased nitrogen metabolism**. In the model, this would be indicated by a decrease in the total velocity of the urea cycle. [14, 15, 27]
3. **Decreased mitochondrial GSH concentration**. In the model, this is assessed through the total glutathione concentration denoted 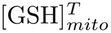 and computed as the time integrated deviation of the metabolite load from baseline. By construction, any nonzero production of H_2_O_2_ yields net GSH depletion through the irreversible detoxification reaction, ensuring 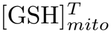 decreases under ethanol perturbation. [9, 10, 28]
4. **Increased fatty acid biosynthesis**. In the model, this would be indicated by an increase in the total velocity of the FABs cycle. [17–19]

In this section, we evaluate whether the model follows such clinically observed phenomena.

#### Model implementation

To assess the above clinically observed attributes, we define two numerical experiments: a simulation where one perturbation of ethanol amounting to 17.4 mM is applied and another where two perturbations of ethanol are applied (72 simulation hours apart) each totaling to 17.4 mM. The ethanol perturbation of 17.4 mM corresponds to a blood alcohol concentration of 0.08 g/dL, the threshold used to define binge drinking [29]. Because the model delivers the dose instantaneously and assumes it is uniformly distributed, the simulated perturbation represents an idealized upper bound. For precise details on the *in silico* experimental setup see Sec. 3.7.

#### Simulation results: validation against clinical observations

The simulations yields the following percentage of cases meeting criterion:

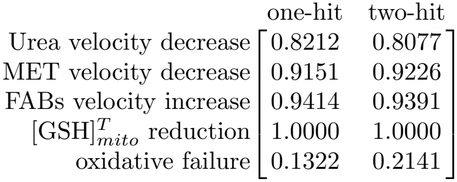

In a vast majority of cases, the model meets the observed physiology, indicating that the four canonical phenotypes emerge from subsystem competition rather than from explicit encoding.

To test whether these phenotypes are a consequence of how ethanol perturbation is encoded, or a genuine result of coarse-graining the biochemical reaction network, we repeat the identical grid search on three altered network topologies (Fig. 2):

1. **Original Network (A):** Network as defined in Sec. 3.4 (Fig. 2.A).
2. **Partial decoupling (B):** Starting from the original network, we remove all weak inter-module interactions while preserving the dominant bidirectional couplings between adjacent cycles. Only the strongest pairwise connections between neighboring modules (MET-GSH, GSH-Urea, Urea-FABs) are retained; all long range and low weight edges are pruned. This yields a locally coupled chain with preserved bidirectional flux between nearest neighbors (Fig. 2.B).
3. **Full decoupling (C):** The network is further reduced by retaining only minimal symmetric interactions within disjoint module pairs. The system is partitioned into two independent subsystems (MET-GSH and Urea-FABs), each preserving only its weakest nonzero bidirectional coupling. All inter-pair connections are removed (Fig. 2.C).
4. **One-way loop (D):** A directed topology is imposed such that all modules remain connected through a single cyclic pathway. Edges form a strongly connected directed loop, ensuring global reachability while eliminating bidirectional redundancy unidirectional flow in contrast to the reciprocal exchange in A and B (Fig. 2.D).

**Fig 2.**
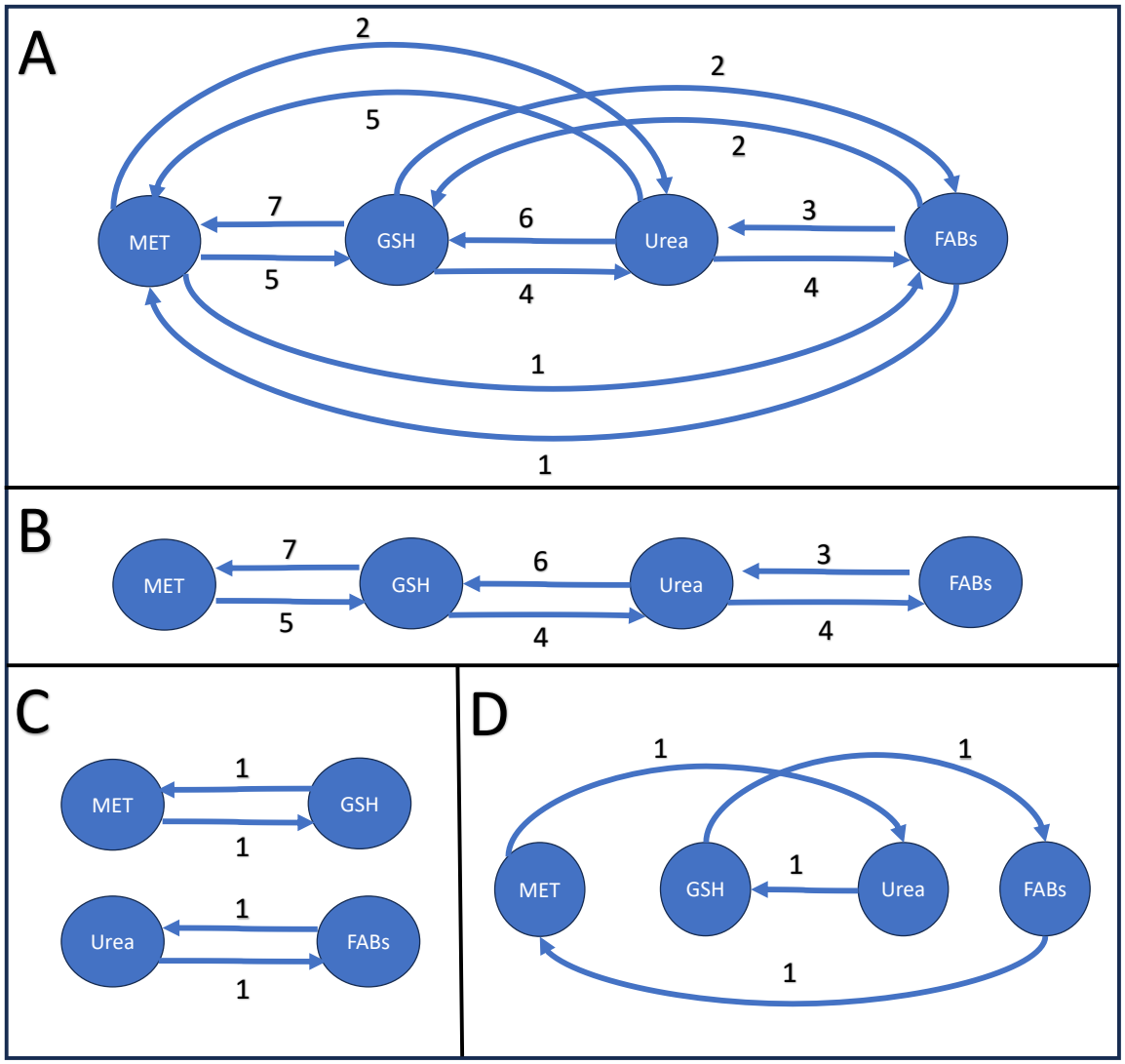
Network diagram connectivity illustrations. (A) Adjacency matrix of the stoichiometrically derived network as defined in Sec. 3.4. (B) Adjacency matrix of the partially decoupled network. (C) Adjacency matrix of the fully decoupled network. (D) Adjacency matrix of the one-way loop network.

Repeating the one-hit grid search across topologies gives the following percentage of cases meeting criterion:

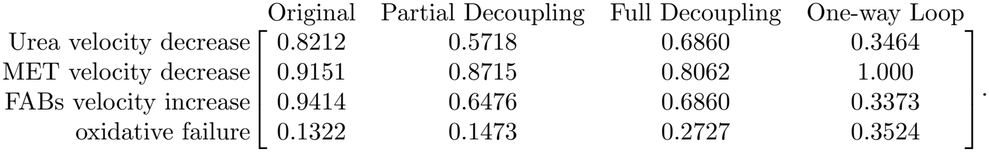

Across all reduced architectures, the fully connected system exhibits the highest agreement with known ethanol-induced phenotypes while maintaining the lowest failure rate. Decreases in urea flux and increases in fatty acid biosynthesis are markedly diminished under partial and full decoupling and largely lost under the one-way loop; by contrast, the decrease in methionine velocity is robust across all configurations, indicating it is driven by local interactions rather than global structure. These results demonstrate that the higher order responses impaired nitrogen metabolism and enhanced lipogenesis emerge from distributed, bidirectional coupling. Disrupting this connectivity both reduces phenotype fidelity and increases instability, indicating that metabolic resilience and phenotype emergence are network level properties requiring reciprocal exchange and redundancy across pathways.

#### Interpretation

The battery of assays demonstrates that the clinically observed phenotypes of ethanol metabolism in the liver can emerge as outcomes of resource competition between coupled metabolic cycles rather than as independent effects. One criterion is an exception worth stating plainly the 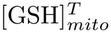 reduction is a direct outcome of the model structure, since any nonzero H_2_O_2_ production forces net GSH depletion, so its 100% rate carries little validating weight. The informative concordances are the urea, MET, and FAB criteria, which the model is free to violate yet satisfies in 80-94% of cases. The topology comparison sharpens the claim: urea suppression and enhanced lipogenesis degrade under decoupling and collapse under unidirectional flow, marking them as properties of reciprocal inter-cycle coupling rather than of how ethanol forcing is encoded, whereas the near invariant MET decrease is locally driven. The significance is that the framework dissociates network level phenotypes from locally determined ones and provides direct evidence that emergent AH behavior is carried by subsystem interlinkage.

### 2.2 Concordance with acute glutathione depletion in vivo

#### Clinical/experimental scenario

We compare the model’s acute GSH response against the mouse ethanol exposure data of Vogt and Richie [30]. They administered 5 g/kg ethanol to C57Bl/6NNIA mice and found hepatic GSH depleted by more than 50% within 6 hours across all age groups, with young and old animals persistently low at 24 hours while mature animals recovered to baseline. Because food was withheld throughout the 24 hour period, the reported trajectories reflect combined ethanol and fasting induced depletion. To isolate the ethanol specific contribution, the authors expressed GSH relative to age matched fasted saline controls, revealing an ethanol attributable depletion of approximately 50-70% at 6 hours. Since the present model does not incorporate fasting dynamics, we benchmark against this ethanol specific signal. The control animals showed no significant fasting induced depletion until after the 6 hour mark, so we validate against the pre-6 hour time points.

#### Model implementation

We adjust the model to mouse metabolism by altering the rate at which ethanol is detoxified from the system. Setting the parameters as defined in Sec. 3.5, we perturb the mouse system with ethanol amounting to 100 mM concentration and compute the percent GSH is depleted from baseline.

#### Simulation results: concordance with percent GSH depletion

At 2 and 6 hours post exposure, [GSH]_*mito*_ is 7.2547 and 4.6127 mM, mirroring the 25% and 51% depletion reported by Vogt and Richie [30]. The model also returns to baseline within the 24 hour window; while this qualitatively matches the recovery of mature mice, the 24 hour measurement in that study reflects combined ethanol and prolonged fasting absent from the present model so this agreement cannot be taken as unambiguous validation at that timepoint. These results are plotted in Fig. 3.

**Fig 3.**
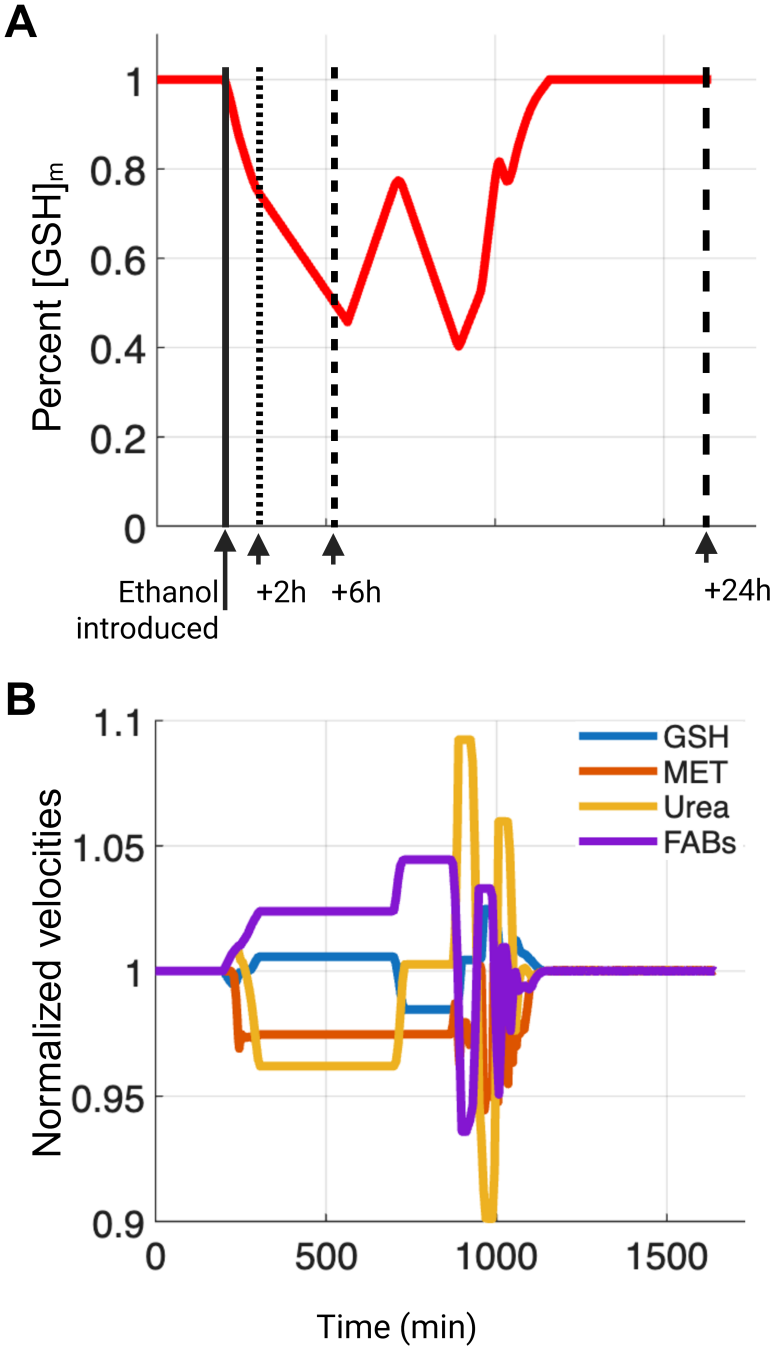
Mouse experiment validation. A simulation tuned to a healthy, adult mouse captures the approximate 25% and 51% depletion of GSH after 2 and 6 hours respectively observed by Vogt and Richie [30]. Note in the plot this is equivalent to the depletion to 75% and 49% after 2 and 6 hours respectively. Solid arrow denotes ethanol introduction to the system, with the increasing size in dashes corresponding to the 2, 6, and 24 hours post ethanol exposure.

#### Interpretation

This model is able to reproduce 25% and 51% depletion at 2 and 6 hours with parameters fixed a priori from physiology, not fit to the trajectory, shows that the framework recovers measured magnitudes, not merely qualitative trends, on a real dataset at the acute timescale that motivates it. Its significance is therefore evidence of calibration free predictive fidelity. The caveats bound the claim tightly: a single species, a single parameter tuning, and the fasting confound that renders the 24 hour recovery uninterpretable, so this validates the depletion limb of the response and not the recovery limb.

### 2.3 Heterogeneous failure analysis uncovers the structure of decompensation

#### Clinical scenario

Patients with comparable ethanol exposure segregate into recoverable and non-recoverable states, exhibiting clinically distinct failure phenotypes. We ask whether the model reproduces this heterogeneity and, if so, what dynamical structure underlies it.

#### Model implementation

To understand the modes of failure, we review in depth four scenarios under 1-hit ethanol perturbations amounting to 17.4 mM. We examine representative healthy, diseased (ALD), failed-compensated (AH), and failed-decompensated (AH) individuals. For further details, see Sec. 3.1.

#### Simulation results: trajectory typology elucidation

In Fig. 4, the plots of [H_2_O_2_]_*mito*_ show that once glutathione buffering is exceeded, mitochondrial H_2_O_2_ accumulates rapidly, with [H_2_O_2_]_*mito*_ ≫ 0 persisting after even a transient excursion above zero. We take this runaway accumulation to define an oxidative failure regime. Concentrations of [H_2_O_2_] above 1 mM are known to be cytotoxic, and we would expect a cascade of cell death within the individual’s liver, denoting an AH scenario [31].

**Fig 4.**
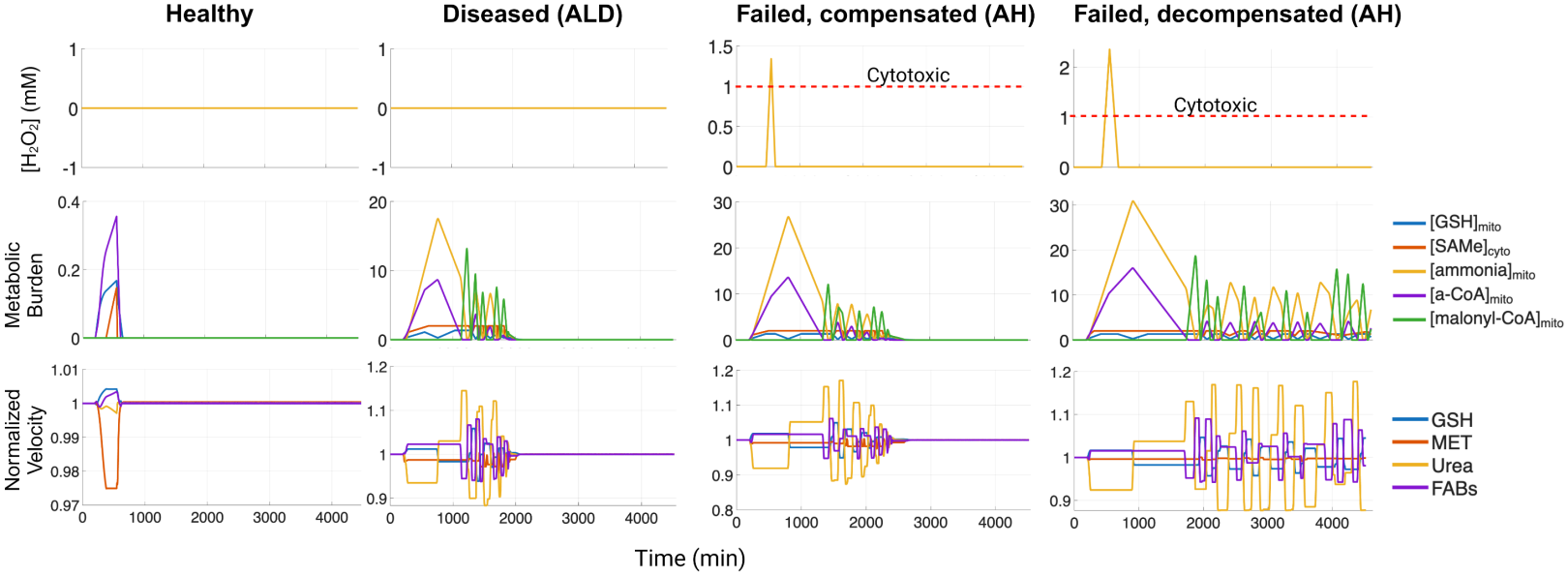
Simulations illustrate four possible states and transitions. The healthy state has very little deviation in any of the three tracked variables from its steady state. The diseased state has a more prolonged and dangerous increase in various metabolite burdens including ammonia. The failed and compensated state has achieves a transient increase in [H_2_O_2_] indicating a cytotoxic state is present. Metabolic compensation is achieved before the end of simulation time and a return to steady state is noted around minute 2500. We define this as nonsevere AH. The failed and decompensated state also has a transient increase in [H_2_O_2_] but does not return to a metabolic steady state. We define this as severe AH.

In Fig. 4, the normalized velocity plots show the modules behaving as coupled oscillators, each velocity responding both to its own state and to its interactions with the others. In the healthy case, perturbations induce small, rapidly damped oscillations and quick relaxation to the initial steady state: a strongly stable regime in which coupling efficiently redistributes transient imbalances. In the diseased and failed states the dynamics change qualitatively, with oscillations growing in amplitude and persistence as damping weakens and coupling turns destabilizing. The failed-compensated regime still returns to baseline, but only after prolonged, higher amplitude oscillations: stability preserved but weakened; the failed-decompensated case shows sustained, uncoordinated oscillations with no rapid relaxation, consistent with a loss of global stability near or beyond a bifurcation threshold. This sustained instability is how we define a metabolically failed regime.

Together, these dynamics support the interpretation of metabolic subsystems as interacting oscillatory units, with disease progression corresponding to a transition from a strongly damped, stable regime to one of amplified oscillations, delayed recovery, and eventual breakdown of coordinated control.

The normalized metabolic burden plots in Fig. 4 mirror this progression. Healthy loads remain tightly regulated with minimal excursions; the diseased state shows transient but recoverable excursions in ammonia and acetyl-CoA; the compensated failure regime shows amplified, prolonged excursions with delayed recovery; and the decompensated regime shows runaway dynamics in which dominant metabolites overwhelm the allocation mechanism and prevent return to baseline. Because the concentration loads directly track the velocity oscillations, this progression is consistent with a critical threshold separating recoverable from non-recoverable states.

#### Model result: two independent failure axes

This abrupt transition marks a critical threshold in the nonlinear dynamics: once effective antioxidant capacity falls below the rate of ROS production, the system cannot return to its pre-perturbation redox equilibrium. The behavior is consistent with a saddle-node type loss of stability in which the stable low H_2_O_2_ branch is annihilated and trajectories diverge toward sustained oxidative accumulation, providing a natural dynamical definition of oxidative failure. A central question is whether metabolic collapse must proceed through this redox bifurcation.

Taking *k*_*d*_ = .001, *y*_1_ = .6, and *y*_2_ = .1 with the same one-hit numerical simulation applied to the example cases above, we can see in Fig. 5 that metabolic decompensation without oxidative failure is possible.

**Fig 5.**
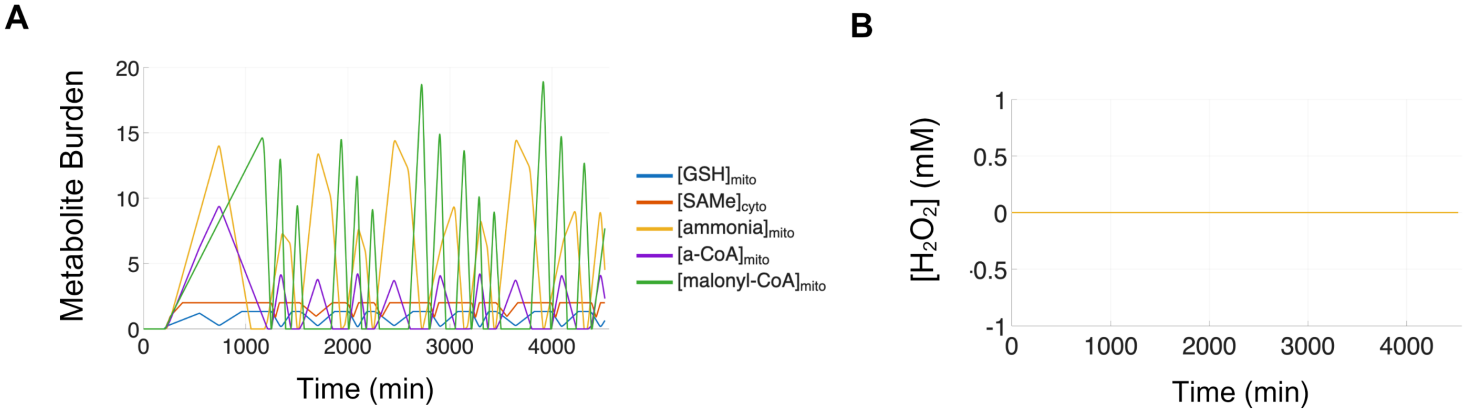
Metabolic decompensation without oxidative failure example. Given *k*_*d*_ = .001, *y*_1_ = .6, and *y*_2_ = .1 with the same one-hit numerical simulation applied to the example cases above, we note that metabolic decompensation can occur without passing through the oxidative failure threshold. **(A)** Metabolic burden is plotted across the simulation and oscialltes without compensating. **(B)** H_2_O_2_ concentration is plotted across the simulation and does not have a transient spike as characteristic of a system undergoing oxidative failure.

Therefore, we can partition long term behavior into two independent mechanisms, loss of oxidative buffering and loss of metabolic flux homeostasis, are formalized as follows:

#### Oxidative failure

Let *T* > 0 denote the terminal simulation time. A trajectory exhibits oxidative failure if there exists *t* ∈ [0, *T* ] such that [H_2_O_2_]_*mito*_(*t*) *> ϵ*_*ox*_ for some *ϵ*_*ox*_ ∈ ℝ^*>*0^ corresponding to the empirically observed redox bifurcation boundary.

#### Metabolic decompensation

Let *t*_*e*_ denote the time at which the final ethanol perturbation has been detoxified, and assume *T* ≫ *t*_*e*_. For fixed constants *δ*_*t*_ ≪ *T* − *t*_*e*_ and *δ*_*i*_ *>* 0, metabolic decompensation occurs if there exists a subsystem *S*_*i*_ ∈ *S* such that

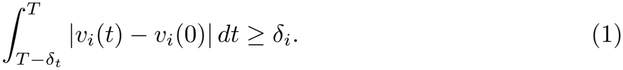

Thus, even after removal of the external perturbation, at least one subsystem fails to relax to its baseline flux.

#### Outcome classification

Let *X* denote the set of admissible initial conditions and parameter realizations, and let *ϕ*|_*a*_ : *X* → *T* map each element to its trajectory given ethanol perturbation series *a*. Define

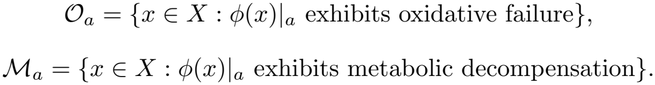

#### Proposition (State space partition into failure regimes)

The trajectory space *X* admits a partition into four disjoint classes, 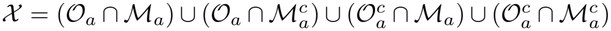, corresponding to regimes with distinct mechanistic and clinical interpretations:

1. **Oxidative failure with metabolic decompensation** (*O*_*a*_ ∩ *M*_*a*_) Oxidative collapse propagates into persistent subsystem flux disruption, producing coupled redox and metabolic failure. Consistent with severe AH, in which inflammatory oxidative injury and metabolic dysfunction coexist.
2. **Oxidative failure without metabolic decompensation** 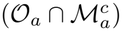. Trajectories cross the redox threshold but recover metabolic flux homeostasis a disease progressive oxidative insult short of full decompensation.
3. **Metabolic decompensation without oxidative failure** 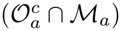. No redox bifurcation is crossed, yet the redistribution network fails to reestablish steady state flux a mechanistically distinct hepatic failure driven by sustained imbalance in ammonia detoxification, methylation, and resource allocation rather than oxidative overload.
4. **Minimal liver disease progression or damage** 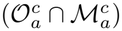. A perturbation inducing neither a bifurcating oxidative insult nor metabolic disruption. We say *minimal* rather than *trivial* because any ethanol perturbation raises FABs and ammonia, which is not biologically benign.

These sets are mutually disjoint and exhaustive (Fig. 6).

**Fig 6.**
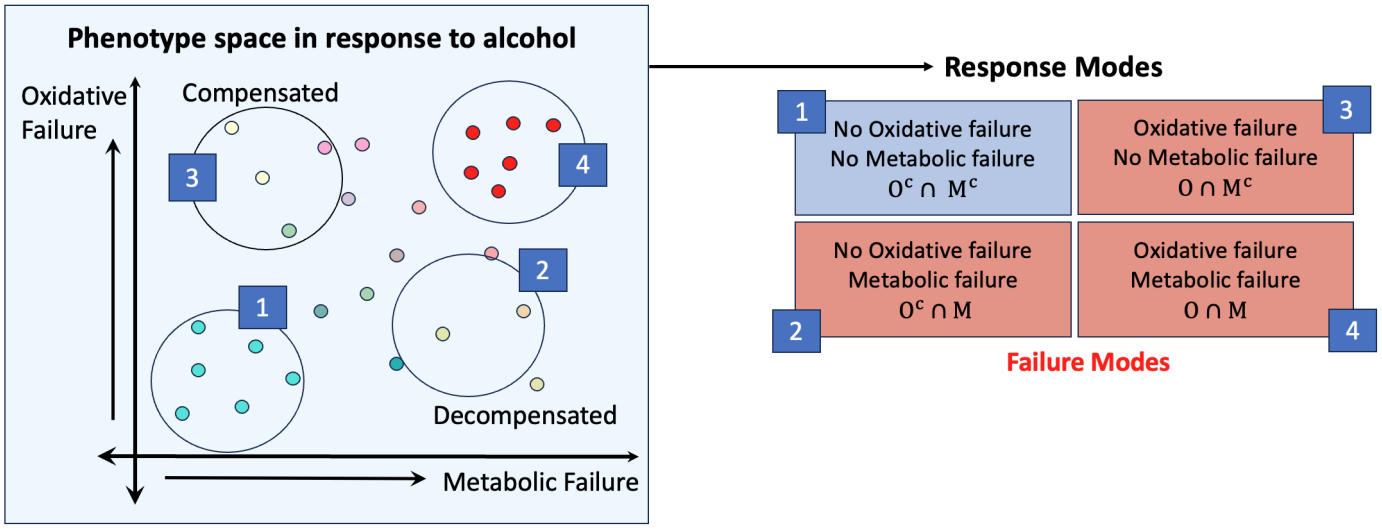
Phenotype to perturbation response: In phenotype space, we have an array of individuals parameterized by *y*_2_, *y*_1_, *k*_*d*_ perturbed with ethanol exposure. This lands these individuals into one of four categories: (1) No oxidative failure and no metabolic failure (2) No oxidative failure but metabolic failure (3) Oxidative failure but no metabolic failure (4) Both o oxidative failure and metabolic failure.

#### Interpretation

The partition reframes decompensation from a single failure mode into two independent axes, oxidative and metabolic failure, that need not co-occur. Clinically, its most consequential prediction is the existence of a purely metabolic failure route 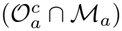 that reaches hepatic failure without ever crossing the redox threshold, a mechanistically distinct phenotype invisible to accounts centered on oxidative stress alone. The saddle-node structure of the H_2_O_2_ transition gives a dynamical basis for the clinical observation that decompensation is abrupt rather than graded: compensation holds up to a critical threshold, beyond which collapse is rapid and irreversible. Casting the subsystems as coupled oscillators supplies a natural vocabulary for severity as progressive loss of damping, connecting a continuous parameter drift to a discontinuous change in outcome.

### 2.4 Cumulative exposure precipitates decompensation

#### Clinical scenario

Repeated drinking episodes, rather than a single exposure, are what clinically precipitate acute-on-chronic decompensation.

#### Model implementation

We compare one-hit and two-hit exposure within the partition of Sec. 2.3.

#### Model result: shift toward joint failure

Applying the partition across the grid we get the following:

#### Percentage of cases in failure

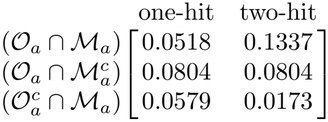

The one-hit system populates all three failure regimes at comparable frequencies, indicating a single perturbation can independently trigger either redox or metabolic instability. The two-hit system shifts pronouncedly toward the joint class (*O*_*a*_ ∩ *M*_*a*_), suggesting repeated perturbations preferentially drive trajectories beyond both recovery thresholds at once. Cumulative alcohol exposure thus induces not merely quantitative worsening but a qualitative transition toward compounded failure.

#### Simulation results: control shifts from parameter driven to interaction dominated

We quantify how patient specific values shape the dynamics using variance based sensitivity analysis on 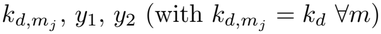, estimating first order and total effect indices via [32] at *N* = 1024 = 2^10^. Two scenarios are compared: a one-hit (17.4 mM) and a two-hit (17.4 mM twice, separated by 240 min).

#### Velocity sensitivity indices (one hit, Fig. 7)

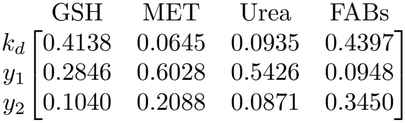

**Fig 7.**
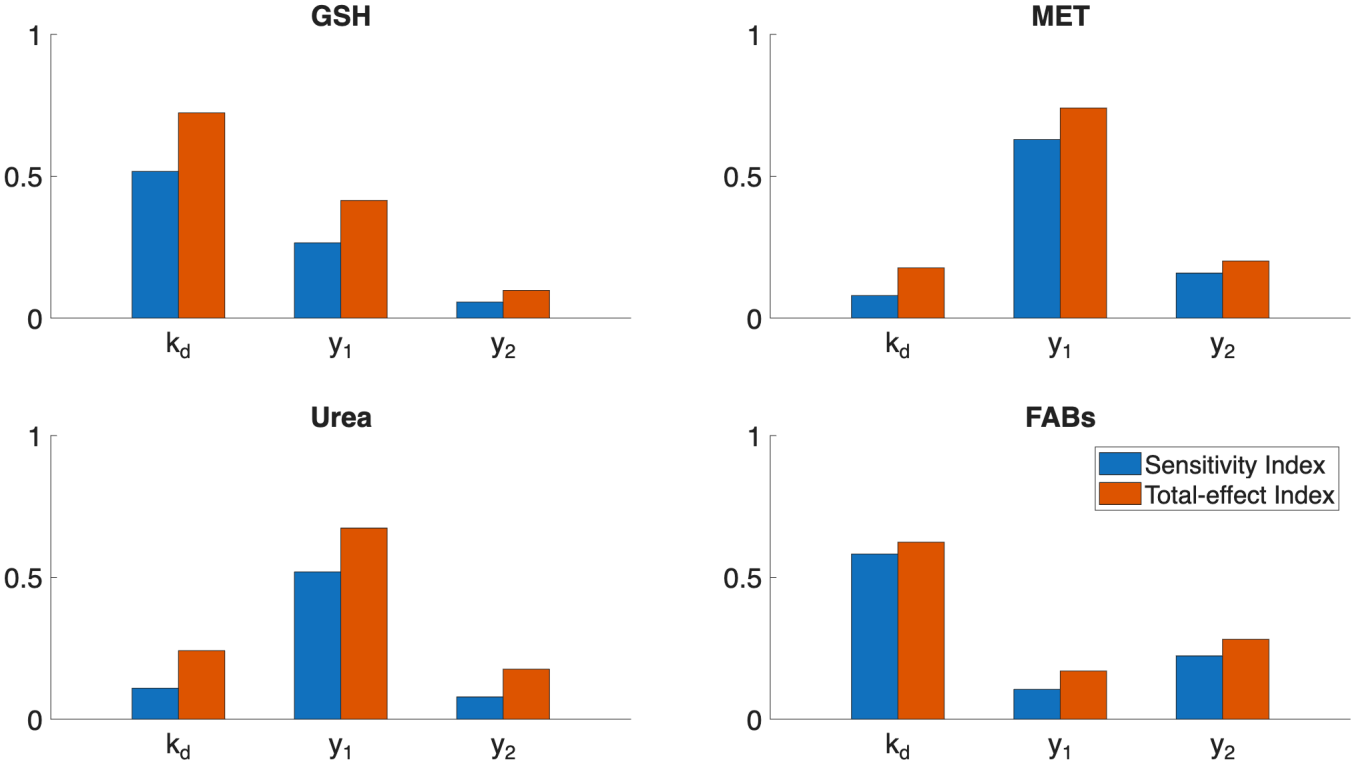
One hit uncertainty quantification reveal patient specific parameter dependencies. Corresponding to each of the three patient specific parameters, both the sensitivity and total effect indices for the velocities of each cycle were computed using variance based sensitivity analysis techniques.

#### Velocity total effect indices (one hit)

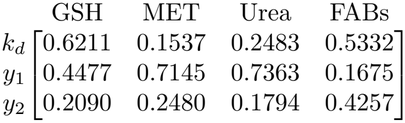

In the one-hit system, first order indices are substantial across multiple subsystems, particularly ROS yield (*y*_1_), which strongly influences MET and urea dynamics; under a single perturbation, key processes retain direct responsiveness to individual inputs.

Total effect indices exceed first order indices everywhere, confirming nonlinear coupling, but the gap is moderate: interactions are present without dominating. GSH remains largely locally governed (primarily by *k*_*d*_), while MET and urea mix direct and interaction driven sensitivity. The one-hit system thus retains partial modularity.

#### Velocity sensitivity indices (two hit, Fig. 8)

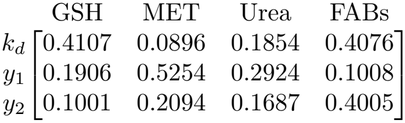

**Fig 8.**
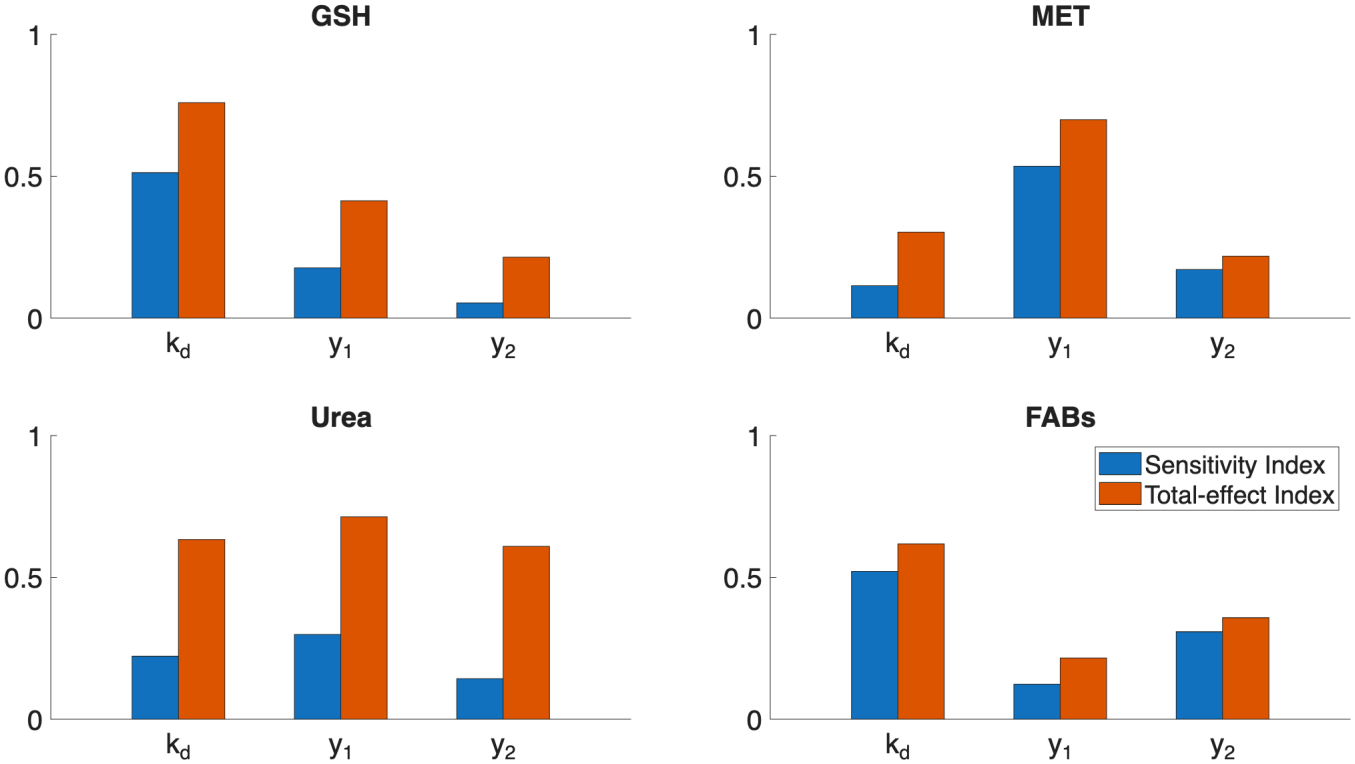
Two hit uncertainty quantification reveal patient specific parameter dependencies. Corresponding to each of the three patient specific parameters, both the sensitivity and total effect indices for the velocities of each cycle were computed using variance based sensitivity analysis techniques.

**Fig 9.**
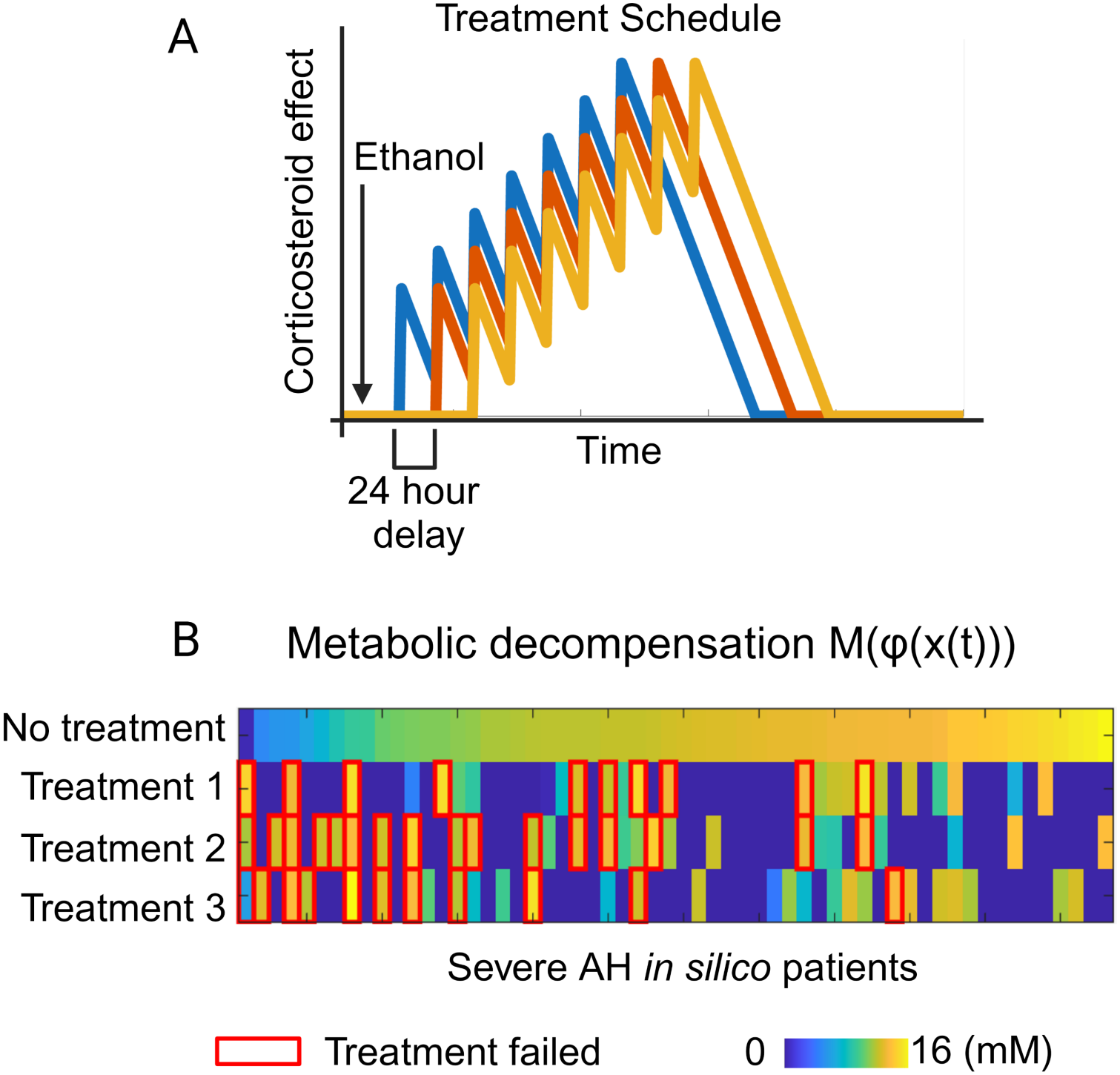
In silico Corticosteroid treatment benchmarks clinically measured success rates: **(A)** Three *in silico* corticosteroid treatment schedules (Blue: 1, Red: 2, and Yellow: 3) were implemented for parameters that resulted in metabolic decompensation after 17 say simulation time (denoting severe AH). **(B)** For each *in silico* severe AH patient, we plot their total metabolic decompensation and indicate whether it defined a treatment success or failure. A treatment regime, *φ*, is deemed successful if the total metabolic decompensation with the treatment is less than that without, i.e. *M* (*φ*(*x*(*t*))) *< M* (*x*(*t*)). Plotted Treatments highlighted in red boxes are those that failed. Treatment was successful in 72 − 82% of the cases, benchmarking the clinically recorded 75% success rate [7].

#### Velocity total-effect indices (two hit)

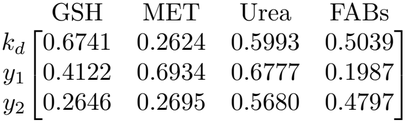

The two-hit system shifts pronouncedly toward interaction dominated dynamics: total effect indices substantially exceed first order indices across all subsystems. The effect is most striking in the urea cycle, where first order sensitivities are minimal while total effects are large, showing that ammonia detoxification is governed almost entirely by global metabolic state rather than any individual parameter. GSH retains some local control (through *k*_*d*_) but with increased interaction contributions; MET and FABs are intermediate but strongly cross coupled; and *y*_1_ exerts large total effects across subsystems, reinforcing its role as a mediator of system wide coupling. This is consistent with the model structure: ethanol metabolism couples directly to GSH through *y*_1_, while acetyl-CoA effects (*y*_2_) distribute across subsystems, and the MET cycle which lacks a direct ethanol yield connection, acquires sensitivity primarily through indirect coupling.

Overall, adding a second perturbation amplifies cross subsystem coupling and reduces the effective independence of metabolic processes. Progression from a single insult to a multi-hit state corresponds to a shift from parameter driven to interaction dominated behavior, consistent with a loss of compensatory capacity and the emergence of system level constraints.

#### Interpretation

The one-hit versus two-hit comparison operationalizes the acute-on-chronic concept: a dose that is individually survivable becomes decompensating when repeated, and the failure distribution shifts specifically toward the coupled class (*O*_*a*_ ∩ *M*_*a*_) rather than worsening uniformly. The uncertainty analysis carries the more refined point: control of the system migrates from parameter driven (locally modular) to interaction dominated (emergent), most starkly in the urea cycle, whose variance becomes governed almost entirely by global state rather than any single input. This is precisely the regime our ammonia hypothesis predicts: once urea cycle behavior is dictated by network state rather than its own local capacity, ammonia clearance can no longer be recovered by that module acting alone, and hyperammonemia emerges as a property of the coupled system rather than of any single failing pathway. The significance is that this provides a quantitative signature of lost compensatory reserve and a direct explanation for why single pathway or single parameter reasoning can succeed early yet fail in advanced disease, where behavior, and the ammonia rise itself, has become a network property.

### 2.5 Clinically observed corticosteroid response heterogeneity is predicted by the model

#### Clinical scenario

Corticosteroids are the mainstay for severe AH, yet roughly 75% of eligible patients respond within 7 days while the remainder do not [7].

#### Model implementation

We ask whether the model is able to capture this heterogeneity. Corticosteroids, which primarily attenuate inflammatory and oxidative processes, are expected to act on trajectories in both *O*_*a*_ and *O*_*a*_. We represent corticosteroid treatment as a reduction in the effective glutathione consumption rate: not a direct elimination of ROS, but a reduction in inflammatory/oxidative burden and thus in the effective demand on mitochondrial GSH buffering. Treatment acts as a crutch for the overall system: the resources that would be utilized by the GSH cycle are then free to be reallocated to other modules.

We take the 72 hour ensemble and identify severe AH trajectories as those in *M*_*a*_, persistently metabolically decompensated at terminal time. We introduce 3 treatment schedules, of which the start time of each is offset by 24 hours. Efficacy of the treatment was evaluated at day 17: by calculating whether or not the system under treatment remained metabolically decompensated. For further details on the numerical treatment schedule see Sec. 3.1.

#### Simulation results: response fraction brackets the clinical benchmark

Of 1331 initial simulations conducted in Sec. 2.1, 209 trajectories fell in *M*_*a*_ under one-hit perturbation. Without treatment, 0.6027 of these regain metabolic stability over 17 days (which we define as non-severe AH); the remaining 0.3973 do not (severe AH). We quantify long-time instability of a trajectory *x*(*t*) through

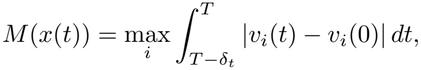

the maximal cumulative terminal window deviation of subsystem velocities from baseline. For a treatment regimen *φ*, we say treatment produces a net stabilizing effect if *M* (*φ*(*x*(*t*))) *< M* (*x*(*t*)). Shasthry et al. reported 75% of corticosteroid eligible severe AH patients classed as responders within 7 days [7]. Assuming all trajectories are eligible, the fraction of severe AH trajectories that under treatment simulation show a positive response ranged from 0.72 to 0.82 across the three schedules, bracketing the clinical benchmark without having been calibrated to it. This provides quantitative support for the resource allocation framework and suggests the model captures the mechanistic basis of corticosteroid efficacy.

Further, of the cases that would otherwise regain metabolic stability, 0.5455 of the treatments would cause the system to lose its metabolic stability. In all, these percentages indicate that treatment outcome depends sensitively on the timing of intervention relative to the system’s oscillatory state. This is in concordance with clinical observations and serves as a mechanistic explanation for this heterogeneity.

#### Interpretation

Reproducing the reported ∼ 75% responder rate within a 0.72-0.82 band, without calibrating to it, is the strongest clinical concordance in the study precisely because the number is an emergent output rather than a fitted target. This reframes steroid non-response as a predictable consequence of a patient’s location in state space rather than an idiosyncratic or dosing failure, and the dependence on intervention window suggests that timing is itself an outcome determinant: both testable, stratification relevant predictions.

### 2.6 Dose and number dependent stability captures a monotone failure threshold

**Clinical scenario**. Decompensation is precipitated in a threshold like manner by cumulative hits. **Model implementation**. We probe the parameter space under multiple perturbations across four scenarios:

1. **Single hit:** 17.4 mM ethanol added.
2. **Double hit:** 17.4 mM added, then a further 17.4 mM after 60 min.
3. **Triple hit:** as double hit, then a further 17.4 mM after 24 h (1440 min).
4. **Quadruple hit:** as triple hit, then a further 17.4 mM after another 60 min. (Fig. 10)

**Fig 10.**
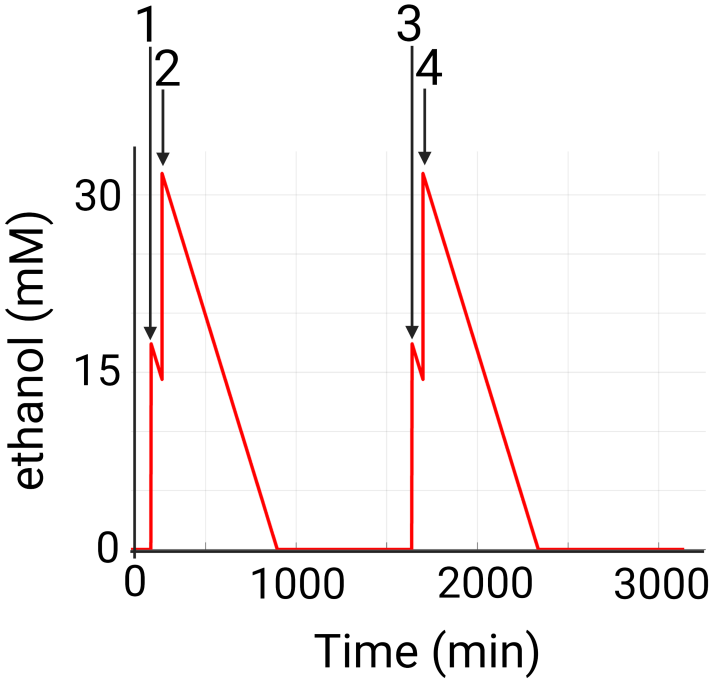
Ethanol quadruple hit dynamics. This plot illustrates how a 4 hit scenario of ethanol ingestion is implemented in the dynamics.

#### Simulation results: monotone failure prediction

We first confirmed that stability under *n* hits implies stability under *n* − 1 hits. We then count how many scenarios a given parameter set withstands without reaching [H_2_O_2_]_*mito*_ *>* 0: in Fig. 11 a three-dimensional phase diagram is sliced through *k*_*d*_, each grid point valued by the number of hits tolerated (a value of 3 meaning stable for the first three but failing the fourth).

**Fig 11.**
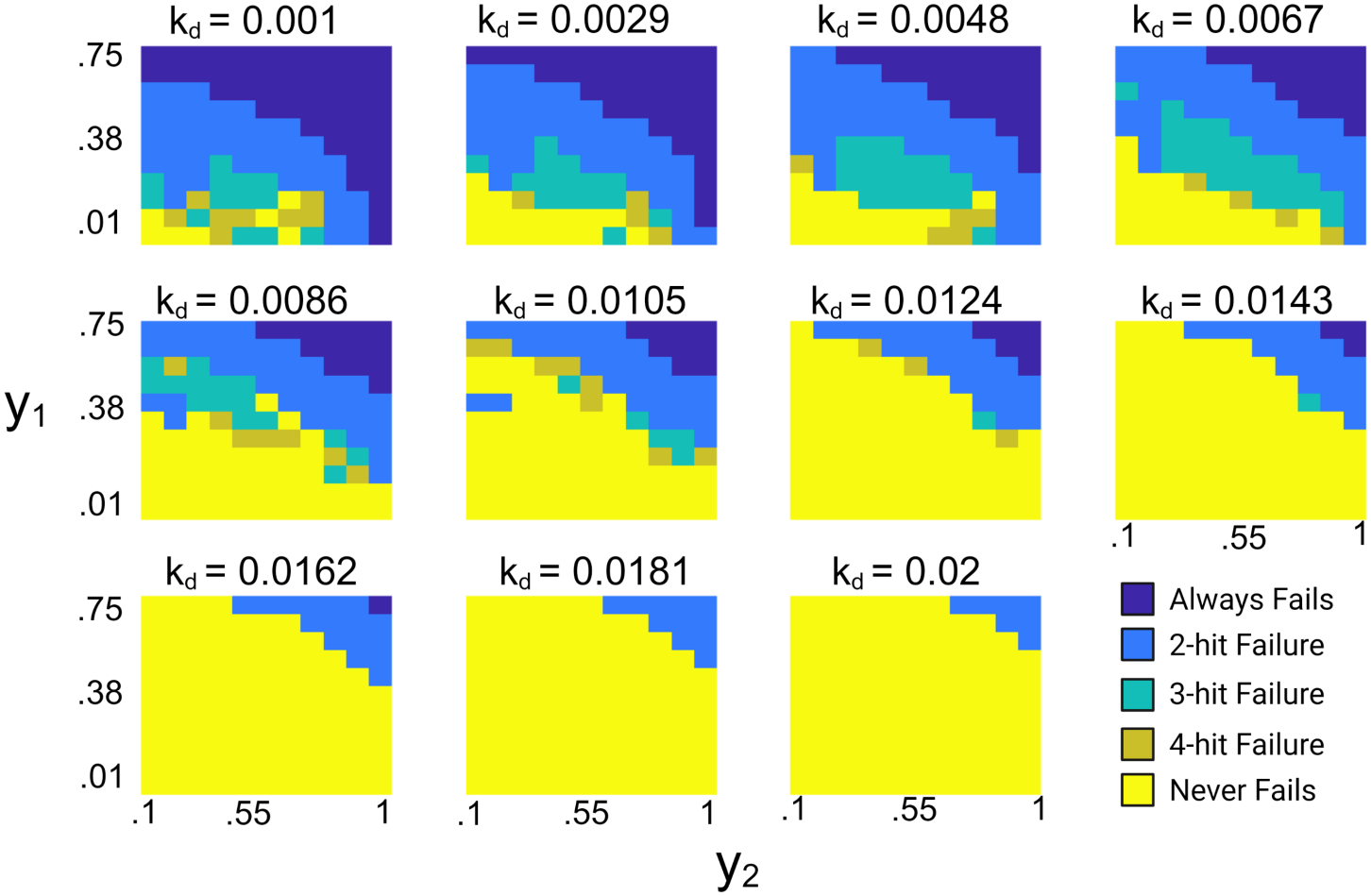
Phase diagram of oxidative stability under multiple hits. Each sub-plot corresponds to a specific *k*_*d*_ value and ranges through (*y*_1_, *y*_2_). Grid point values give the number of ethanol perturbations tolerated without reaching [H_2_O_2_]_*mito*_ *>* 0; a value of 3 is stable for the first three scenarios but fails the fourth.

The stability landscapes across (*y*_1_, *y*_2_) depend strongly on *k*_*d*_, which acts as a global control parameter for robustness. At low *k*_*d*_, stability is confined to a small region and most configurations fail under repeated perturbation. As *k*_*d*_ increases, a sharp transition boundary emerges and propagates across the domain, separating stable and unstable regimes along a diagonal *y*_1_, *y*_2_ trade-off. Intermediate *k*_*d*_ shows coexisting regions of differing stable case counts (0-4), a heterogeneous, threshold dominated response consistent with bifurcation. At sufficiently high *k*_*d*_ the system is nearly uniformly stable, showing that increased decay mitigates accumulation driven instability. Because the second and fourth perturbations arrive before the first and third are fully metabolized, any set that withstands the first but not the second (or the third but not the fourth) delineates a dose dependent domain: the system bifurcates not only across parameter space and hit number but also with respect to ethanol concentration.

#### Interpretation

The phase diagrams (Fig. 11) identify the transport coefficient *k*_*d*_, effective metabolite permeability and clearance, as a single, interpretable global control on robustness, collapsing a high dimensional stability question onto one physiologically meaningful axis. Its significance is twofold: it offers a candidate stratification variable, with robustness set by clearance capacity, and it implies a therapeutic concept distinct from oxidative burden reduction, namely restoration of transport and clearance. The joint bifurcation across both hit number and ethanol concentration formalizes the clinical intuition of a point of no return, and the nesting property (stability under *n* hits implying stability under *n* − 1) endows the dose dependence with a clean monotone structure rather than an erratic one.

## Discussion and Conclusions

We have presented a reduced order, flux-centric framework for studying metabolic reprogramming in alcohol-associated liver disease, built on the premise that hepatic metabolism can be coarse grained into a small number of interacting subsystems competing for a shared, finite metabolic resource budget. Rather than resolving detailed kinetic equations, the model tracks how metabolic flux is redistributed among the glutathione, methionine, urea, and fatty acid biosynthesis cycles under ethanol perturbation. The model simulations demonstrate that multiple clinical phenotypes of ethanol metabolism in ALD can emerge from the interlinkage of these metabolic subsystems (Sec. 2.1). The fully connected network reproduces these phenotypes with the highest fidelity and lowest failure rate, while decoupled and unidirectional alternatives do not, indicating that ethanol response phenotype emerges as a network level property requiring reciprocal exchange across pathways.

Beyond reproducing known physiological/clinical phenomena, the modeling framework characterizes ALD progression as a bifurcation-like loss of stability rather than a gradual degradation across stages. In this framework, metabolic cycles behave as coupled oscillators whose damping weakens as parameters shift toward disease regimes. Failure is signaled by runaway mitochondrial H_2_O_2_ and occurs abruptly once buffering capacity is exceeded, consistent with a saddle-node transition (Sec. 2.3). Critically, this transition is perturbation-history dependent such that an ethanol intake that is episodically tolerated can precipitate collapse when repeated over a compressed timeline. The same cumulative dose produces qualitatively different outcomes depending on its temporal structure (Secs. 2.4 and 2.6). Our modeling framework recasts acute-on-chronic decompensation as a dynamical threshold phenomenon rather than the endpoint of linear accumulation.

Our modeling results are consistent with clinical literature on AH characteristics and outcomes. Primarily, The model offers a mechanistic account of hyperammonemia that does not require epigenetic changes within the context of the urea cycle. Elevated blood ammonia is a robust marker of severity and mortality in severe AH [14, 33, 34], and urea synthesis is known to decrease after ethanol exposure even in healthy individuals [15], yet the canonical oxidation intermediates generate no ammonia [27, 35]. In our framework the suppression of the urea cycle arises upstream in the interconnected metabolic network. Under oxidative stress, glutathione synthesis diverts glutamate away from *N* -acetylglutamate production, diminishing the obligate allosteric activation of CPS1 [36, 37] and throttling the urea cycle. Hyperammonemia thus emerges as a consequence of resource competition between the GSH and urea subsystems rather than as a direct effect of ethanol metabolism (Sec. 2.1). Sensitivity analysis of the two-hit scenario indicated that the separation of the urea cycle is governed by a global metabolic state rather than the cycle specific parameters alone.

Second, the framework assigns mechanistic weight to both the dose and the timing of ethanol intake episodes. The one-hit versus two-hit comparison shows that the failure distribution shifts specifically towards the coupled oxidative and metabolic modes. The sensitivity analysis revealed the flux-control shifted from parameter driven to interaction dominated as a second perturbation is added in the two-hit scenario (Sec. 2.4). The multi-hit phase diagrams extend this result, exhibiting bifurcation not only across parameter space and number of hits but also with respect to whether successive doses arrive before the previous one is fully cleared (Sec. 2.6). These outcomes are consistent with the clinical phenomena in which cumulative and closely spaced ethanol exposures precipitate liver metabolic decompensation [38, 39]. And these results imply that drinking pattern, not merely total intake in a given time-span, is mechanistically relevant to acute-on-chronic risk of liver failure. It also formalizes why single pathway or single parameter reasoning may explain early disease phenomena yet fail in accounting for advanced disease characteristics, where the liver response emerges from distributed control across multiple subsystems. Further, by modeling treatment as a reduction in oxidative demand on the glutathione subsystem, the framework reproduces the reported responder fraction without explicit calibrated (Sec. 2.5) [6, 7].

The framework rests on a small number of defining features. It is flux-centric. The dynamical state variables are the effective velocities of coarse-grained metabolic modules rather than metabolite concentrations, so the model resolves how the network reallocates its activity under stress rather than how individual concentrations evolve. Additionally, our model formulation imposes a conservation constraint drawn from metabolic control analysis. The total velocity across modules is fixed over the acute timescale, the sum of steady-state capacity is conserved in analogy to flux control coefficients, and perturbation can therefore only redistribute capacity, never create it. Metabolite variables enter not as concentrations but as load signals, analogous to elasticities, that quantify the demand or accumulation pressure a deviation from baseline places on their corresponding metabolic cycles. All coupling is derived from stoichiometry, constructed directly from the reactions of the genome-scale reconstruction Human1, so inter-module interactions reflect shared reactants and cofactors rather than fitted interaction strengths. The model is minimally parameterized: baseline module velocities are inferred from patient liver transcriptomes, and individual variation is captured by only three patient-specific coefficients, the ROS yield *y*_1_, the acetyl-CoA yield *y*_2_, and the transport coefficient *k*_*d*_. Together, these features sidestep the kinetic parameters that are unidentifiable at organ scale while ensuring that pathway-level phenotypes arise from competition among modules rather than being encoded directly.

Several limitations follow from these choices. The framework assumes a fixed total resource budget over the acute timescale and therefore does not capture longer term adaptation through gene regulation or protein synthesis. Furthermore, framework parameters are drawn from physiological ranges and proportionalities rather than from direct measurement. Validation is correspondingly qualitative, ordinal, and based on emergent dynamics rather than a calibrated fit to quantitative data. A key limitation is the lack of hours scale longitudinal intrahepatic metabolomics from AH patients. In addition, patient specific parameters that set individual dynamics (*y*_1_, *y*_2_, and *k*_*d*_) cannot be measured directly at present. These are field-level gaps rather than deficiencies unique to the model, and they restrict the model tuning and subsequent qualitative and quantitative validation.

The natural path forward to addresses the above listed limitations would be to acquire patient specific longitudinal data to enable personalized parameter estimation and convert the present qualitative validation into a calibrated one. Further, allowing the total resource capacity to adapt would extend the framework from the acute-on-chronic window to the chronic progression that precedes it. Coupling this reduced order description to genome scale flux models would yield a multiscale approach linking mechanistic biochemical detail to the emergent, system level dynamics the framework was built to capture.

Although our modeling framework is implemented to model the decompensation associated with alcohol-associated hepatitis, the general formalism is neither specific to this disease nor to the liver itself. The model inputs are tissue-agnostic: metabolic cycles are constructed from an atlas of the human metabolism, baseline velocities can be inferred from any tissue’s transcriptome, and the transport coefficient *k*_*d*_ already represents exchange across compartmental boundaries. Therefore, it can be applied to a wide range of acute-on-chronic disorders which are characterized by a gradual erosion of coupled metabolic systems followed by an acute and sudden shift in compensation. As an acute-on-chronic illness, type 1 diabetes illustrates the possible extensions of our modeling framework. Autoimmune *β*-cell loss progresses silently through presymptomatic stages before clinical onset, frequently as diabetic ketoacidosis.

Diabetic ketoacidosis is a medical emergency that if left untreated results in death. This onset threshold reflects the balance between remaining insulin-production capacity and insulin demand rather than *β*-cell loss alone [40], mirroring the capacity-demand structure of the present framework [41].

## 3 Materials and methods

### 3.1 Biological components

Our framework coarse-grains hepatic metabolism into four interacting metabolic modules: the glutathione (GSH) cycle, the methionine (MET) cycle, the urea cycle, and even-chain fatty acid biosynthesis (FABs).

Each subsystem is chosen for its distinct homeostatic role related to ethanol metabolism. The glutathione cycle is the primary cellular defense against mitochondrial reactive oxygen species (ROS), so that maintenance of adequate mitochondrial GSH is essential to prevent oxidative damage, which is a canonical feature of ethanol metabolism [42]. The methionine cycle maintains cellular methylation capacity and regulates gene expression through S-adenosyl-L-methionine (SAMe), the principal biological methyl donor, which additionally acts as a regulator of GSH synthesis [11, 13]. The urea cycle is the primary route of ammonia clearance, and its dysregulation contributes to the elevated blood ammonia that is a hallmark of AH severity [14, 27]. Finally, even-chain fatty acid biosynthesis serves as the principal mitochondrial acetyl-CoA sink, a key driver of hepatic steatosis [17–19].

The effect of ethanol on each of these four metabolic modules is experimentally well characterized. Ethanol is oxidized in the liver via three primary pathways alcohol dehydrogenase (ADH), catalase, and cytochrome P450 2E1 (CYP2E1) [35]. Under normal physiological conditions the majority of ethanol is metabolized via ADH, and the resulting excess NADH is reoxidized through the mitochondrial electron transport chain, particularly at Complex I, a process that generates ROS [43]. Under chronic alcohol exposure or acute high dose (binge) intake, ethanol metabolism via CYP2E1 is upregulated [1, 28, 35]; because cytochrome P450 enzymes are prone to uncoupling, this pathway leaks additional ROS, including H_2_O_2_ [44]. Mitochondrial H_2_O_2_ is detoxified predominantly by the glutathione peroxidase pathway [42], placing a direct oxidative load on the GSH cycle. In parallel, ethanol oxidation proceeds through acetaldehyde to acetate and ultimately to mitochondrial acetyl-CoA, accumulation of which promotes fatty acid biosynthesis [17–19]. Ethanol exposure also disrupts methylation capacity through the methionine cycle [13], and suppresses urea synthesis even in otherwise healthy individuals [14, 15]. In summary, ethanol metabolism in the liver is marked by an increase in ROS that is detoxified by GSH, a decrease in methylation potential, a decrease in nitrogen metabolism, and an increase in fatty acid biosynthesis.

The suppression of urea synthesis is, at first glance, paradoxical, and resolving it motivates the central coupling in our model. Elevated blood ammonia is consistently observed as a strong indicator of disease severity and mortality in AH, despite ammonia not being a direct byproduct of ethanol metabolism [27, 35, 45]. The urea cycle, the primary route of ammonia detoxification, is rate limited by carbamoyl phosphate synthetase I (CPS1) [36], whose activity is strongly dependent on its essential allosteric activator *N* -acetylglutamate (NAG) [37]. NAG is synthesized from glutamate and acetyl-CoA; because ethanol metabolism elevates acetyl-CoA, a purely kinetic argument would predict increased NAG production, and thus enhance urea flux the opposite of what is observed. This implies that the other reactant, glutamate, must be rate limiting this cycle. Glutamate is also the key precursor for glutathione synthesis. Hence under ethanol induced oxidative stress, it is likely that the elevated demand for mitochondrial GSH imposes a dominant sink on glutamate, so that glutathione synthesis outcompetes the urea cycle for glutamate, leading to reduced NAG production, diminished CPS1 activation, and ultimately impaired ammonia detoxification. We therefore propose that hyperammonemia emerges not as a direct consequence of ethanol metabolism but as an emergent property of competition between the GSH and urea subsystems for a shared resource the coupling that our model aims to make explicit. Further, we will examine whether urea synthesis inhibition under ethanol metabolism can be explained independent of gene regulatory actions that could decrease the efficacy of the urea cycle.

We consider these four metabolic cycles as they constitute the dominant flux “handles” governing the metabolites produced or disrupted by ethanol metabolism, and they are interdependent through shared resources: glutamate, acetyl-CoA, and NAD^+^/NADH redox carriers. Consequently, the failure of one subsystem directly imposes metabolic burden on the others. We postulate that the observed systems level decompensation in AH arises from this competition for shared metabolic resources.

### 3.2 Mathematical formalism

We consider metabolic *modules* (or subsystems) as the primary dynamic entities. Each module represents a coarse-grained metabolic pathway whose activity level can adapt in response to metabolic perturbations.

Let *S* = *{S*_1_, …, *S*_*N*_ *}* denote the set of metabolic modules. Each module, *S*_*i*_, corresponds to an aggregated metabolic pathway (for example the glutathione cycle or the urea cycle) with an effective velocity, 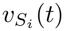, representing the net rate at which that module processes metabolites after accounting for regulatory interactions and resource availability.

We consider a set of metabolites critical to the modules of interest, *M* = *{m*_1_, …, *m*_*N*_ *}*, and associate a time dependent variable, [*m*_*j*_](*t*), to each *m*_*j*_ ∈ *M*. The equation [*m*_*j*_](*t*) represents an effective metabolite load and serves as a coarse-grained control variable linking metabolite level to the resource allocation structure of the module network.

Modules interact with multiple metabolites that are either consumed or produced, and conversely a single metabolite may be processed by multiple modules. We encode this relationship through a coupling matrix

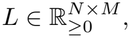

where 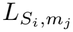 denotes the amount of metabolite *m*_*j*_ consumed or produced per unit velocity of module *S*_*i*_. This is determined through stoichiometry of the biochemical reactions that compose the metabolic module *S*_*i*_.

#### 3.2.1 Constructing metabolic modules

To construct metabolic modules from a detailed reaction network, we begin with a pathway of interest. Let *L*_1_(*S*_*i*_) denote the set of reactions directly associated with the module *S*_*i*_. We then expand this set by including reactions that produce the reactants used in *L*_1_. This expanded set of reactions is denoted as *L*_2_(*S*_*i*_), where *L*_1_(*S*_*i*_) ⊆ *L*_2_(*S*_*i*_).

Modules constructed in this manner are not assumed to be isolated. Instead, they interact through shared metabolites and cofactors. These interactions define a directed network between modules, represented by an adjacency matrix

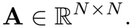

The entry 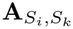 measures the extent to which module *S*_*i*_ can utilize resources that are also required by module *S*_*k*_. Formally,

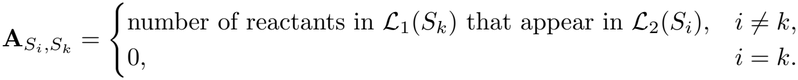

Because metabolic reactions are not always reversible, this interaction network need not be symmetric. See Fig. 12 for a depiction of the generalized formalism.

**Fig 12.**
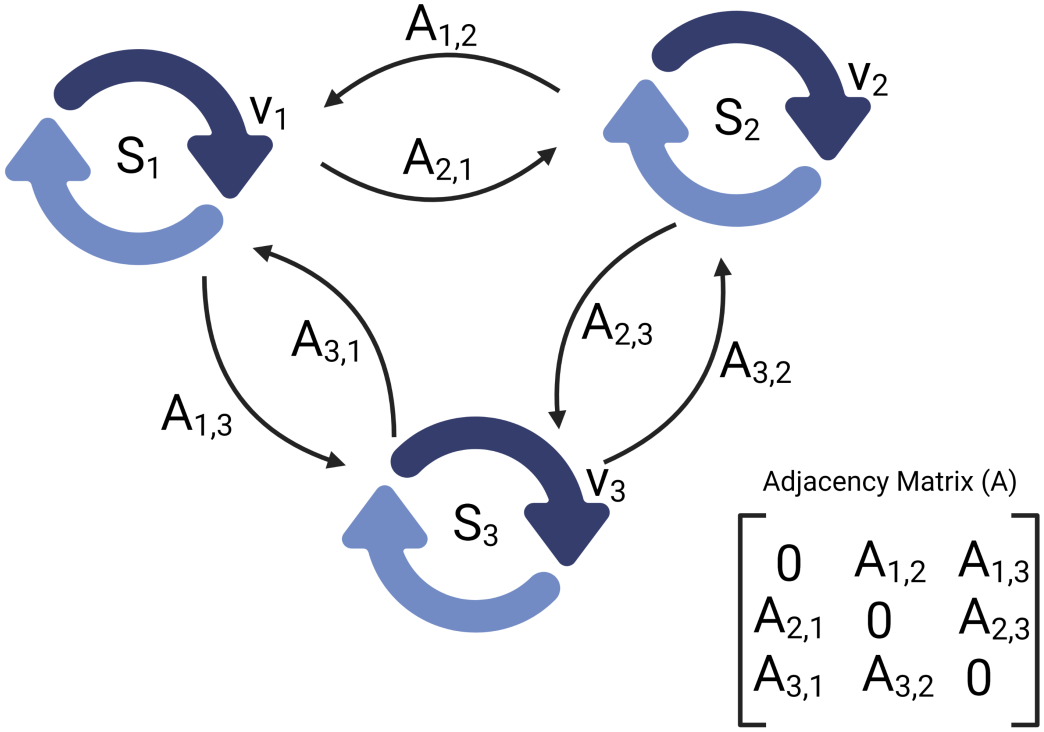
Depiction of generalized formalism links between metabolic modules: We construct each metabolic module *S*_*i*_ by beginning with a pathway of interest. We then expand this set by including reactions that produce the reactants used in the original set of reactions. We construct the adjacency matrix, *A*, such that entry 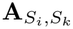 measures the extent to which module *S*_*i*_ can utilize resources that are also required by module *S*_*k*_.

#### 3.2.2 Metabolite dynamics

Metabolites are partitioned into two classes:

(Class 1) metabolites for which the critical accumulation threshold lies closer to the physiological range than the critical depletion threshold, and (Class 2) metabolites for which the critical depletion threshold lies closer to the physiological range than the critical accumulation threshold.

At each point in time, the system reallocates resources across metabolic modules in response to the imbalances arising from demand–supply interactions. In the following, we first define the dynamics associated with each metabolite class independently, and then describe how these contributions are integrated to recover the full module level dynamics.

#### Class 1: Critically high dynamics

We first consider metabolites for which the critical accumulation threshold lies closer to the physiological range than the critical depletion threshold. These metabolites are therefore primarily constrained by the risk of accumulation and must be maintained through continuous consumption. A canonical example is ammonia within the urea cycle. Let *m*_*j*_ denote such a metabolite, and, without loss of generality, shift the reference frame so that its steady state level corresponds to [*m*_*j*_] = 0.

Each metabolite *m*_*j*_ is produced at a baseline rate 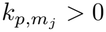 by processes external to the module responsible for its regulation. Under unperturbed conditions, the associated metabolic module operates at an effective capacity 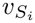, where 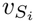 denotes the dynamically allocated metabolic velocity. Deviations from steady state induce a redistribution of this capacity through the resource allocation mechanism. In addition, the parameter 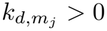 represents an effective transport flux that captures the system’s ability to restore metabolite balance through exchange with the surrounding environment. This includes both removal of excess metabolite from the compartment and replenishment under depletion. Accordingly 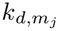 should be interpreted as a coarse-grained representation of aggregate transport processes rather than a single mechanistic pathway. Biologically, this parameter reflects the effective permeability of the cellular environment, quantifying how efficiently the system maintains metabolic homeostasis through exchange across compartmental boundaries.

Consistent with the modeling framework, these terms should be interpreted as contributing to the effective load on the system rather than as components of a detailed kinetic mass balance.

All rates satisfy

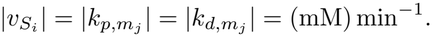

The total clearance velocity of metabolite *m*_*j*_ is then defined as 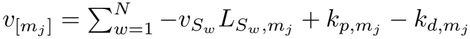. We can then define the metabolite dynamics as

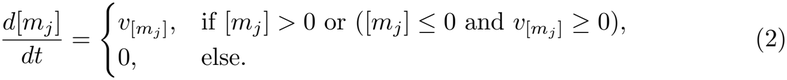

This is linear decay with a hard non-negativity constraint reflects the cessation of detoxification once the metabolite is cleared. Consistent with the flux-centric perspective of the model, this equation should not be interpreted as a detailed kinetic mass balance, but rather as a coarse-grained representation of how metabolite load evolves under net system level fluxes. Note that 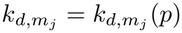 is a patient specific parameter whose estimation is detailed in Sec. 3.4.

Following an acute perturbation that elevates metabolite levels, modules must redistribute limited metabolic resources. At steady state the total processing rate for metabolite *m* is:

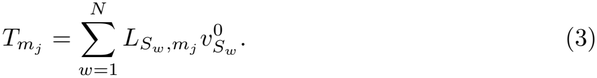

The fraction of metabolite processing attributed to each module is therefore

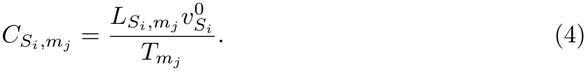

Note that this value is not time dependent and represents a total capacity on the system to metabolize *m*_*j*_ at steady state. Let Δ*t* denote the characteristic clearance timescale. If metabolite concentration increases by Δ*m*_*j*_, the *demand* placed on module *S*_*i*_ by metabolite *m*_*j*_ is approximated by

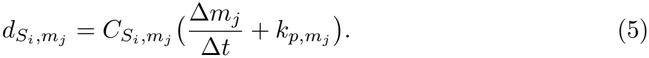

Let 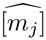 denote the concentration beyond which metabolite *m*_*j*_ causes cellular damage and 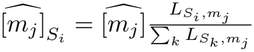 be the fraction of criticality of module *m*_*j*_ attributed to cycle *S*_*i*_. The fraction of resources available for redistribution is

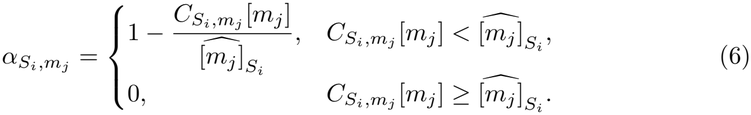

We also define the normalized burden exerted on metabolic module *S*_*i*_ by metabolite *m*_*j*_ as

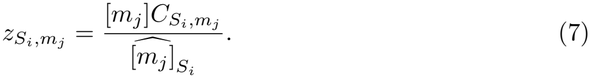

#### Class 2: Critically low dynamics

We now consider metabolites whose depletion is harmful and must be restored by metabolic production modules, such as S-adenosylmethionine (SAMe) in the methionine cycle. Let *m*_*j*_ denote such a metabolite and let *S*_*i*_ denote the module primarily responsible for maintaining its level.

We assume metabolite *m*_*j*_ is consumed externally at baseline rate 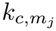 by processes outside of *S*_*i*_, while the module operates at an effective production velocity 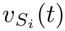. Additionally, recall the parameter 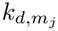 which represents the system’s ability to restore metabolite balance through transport processes that remove excess metabolites from the compartment or import depleted metabolites into it.

To prevent unbounded accumulation, we impose an admissible upper bound 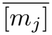. Let 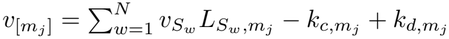. The produced metabolite dynamics are

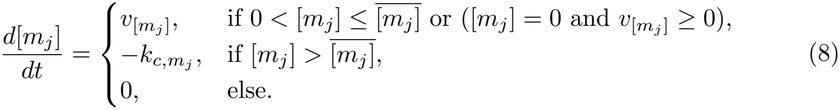

Thus, when the metabolite is above its admissible range, net production is shut off and external consumption drives it back toward the admissible regime; when the metabolite reaches zero, production cannot drive it negative.

Following an acute perturbation that elevates metabolite levels, modules must redistribute limited metabolic resources. At steady state the total production rate for metabolite *m* is again:

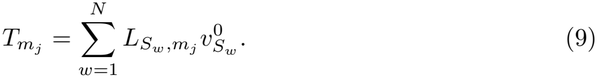

The fraction of metabolite processing attributed to each module is therefore

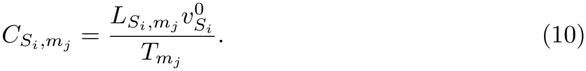

Following an acute depletion of *m*_*j*_, production modules must reallocate limited shared resources to restore metabolite levels. Just like before, Δ*t* denotes a characteristic restoration timescale. The instantaneous demand placed on module *S*_*i*_ by metabolite *m*_*j*_ is

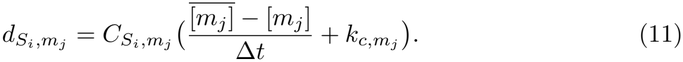

Let 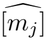 denote a critical lower bound below which depletion causes damage, and let 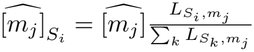 be the fraction of criticality of module *m*_*j*_ attributed to cycle *S*_*i*_. The fraction of allocatable resources that module *S*_*i*_ in terms of metabolite, *m*_*j*_, can share is

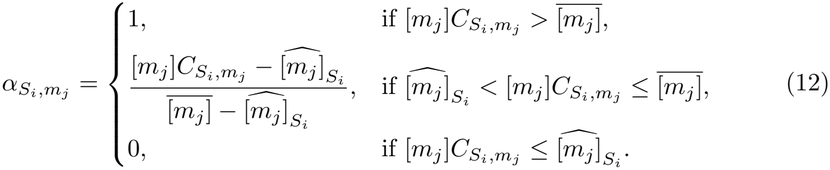

To prioritize restoration under severe depletion, we define a normalized depletion burden of metabolic module *S*_*i*_ with respect to metabolite *m*_*j*_

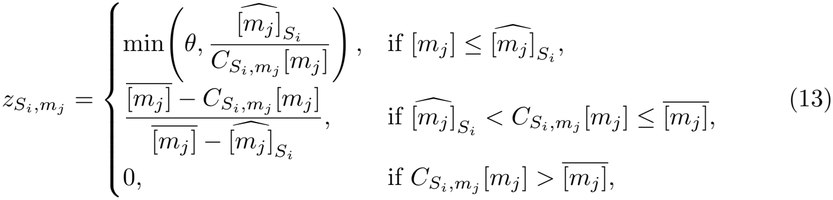

where *θ* is a large bounded constant to avoid singularity as [*m*_*j*_ ] → 0^+^.

#### Metabolic module dynamics

To redistribute resources, we aggregate the metabolite-level quantities above to the level of modules. For module *S*_*i*_, let 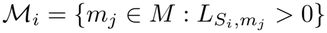 denote the metabolites it produces or consumes. The demand on *S*_*i*_ is set by its most demanding metabolite,

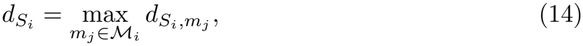

and the fraction of its activity it can spare is set by its most constrained metabolite,

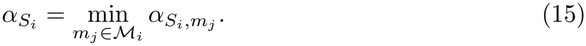

The shareable pool of *S*_*i*_ is therefore

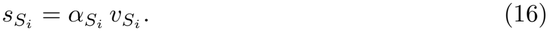

Similarly, the normalized burden on *S*_*i*_ is

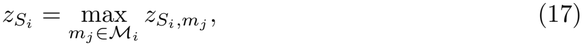

and modules are prioritized through the weights

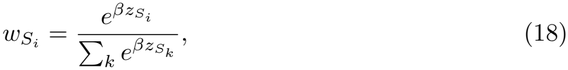

where *β >* 0 controls prioritization sharpness. This choice gives smooth prioritization while allowing sharp switching as *β* increases.

A module whose demand exceeds its current velocity has an unmet demand

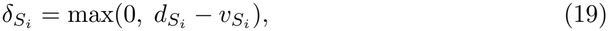

which produces the request intensity

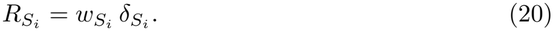

Requests are routed across the module interaction network in proportion to shared resources,

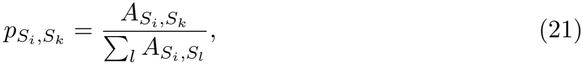

and each donor *S*_*k*_ divides its shareable pool among the modules requesting from it, in proportion to their routed requests,

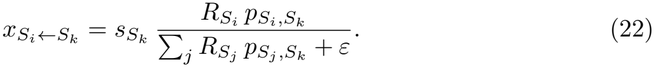

Module velocities then change in proportion to their net transfer,

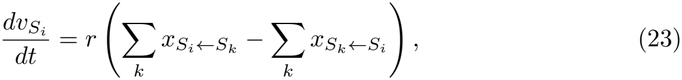

where *r >* 0 (min^−1^) is the response rate. Because each transfer leaves one module and enters an other, summing Eq. (23) over modules gives 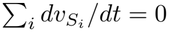. The total velocity 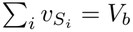 is therefore conserved exactly at every instant: perturbation cannot create capacity, only move it between modules.

Equation (23) closes the coupled system. Metabolite loads set each module’s demand, availability, and burden. These determine the transfers, the transfers change the velocities, and the velocities in turn set how quickly each load is cleared or restored. Each velocity obeys a first-order equation. The oscillatory behavior therefore does not come from inertia in any single velocity. It arises from the feedback loop between velocities and metabolite loads: a module that gains capacity clears its metabolite, which lowers its burden and demand and releases capacity to other modules, whose loads then rise in turn. An example of these dynamics for a two module, two metabolite system can be seen in Fig. 13.

**Fig 13.**
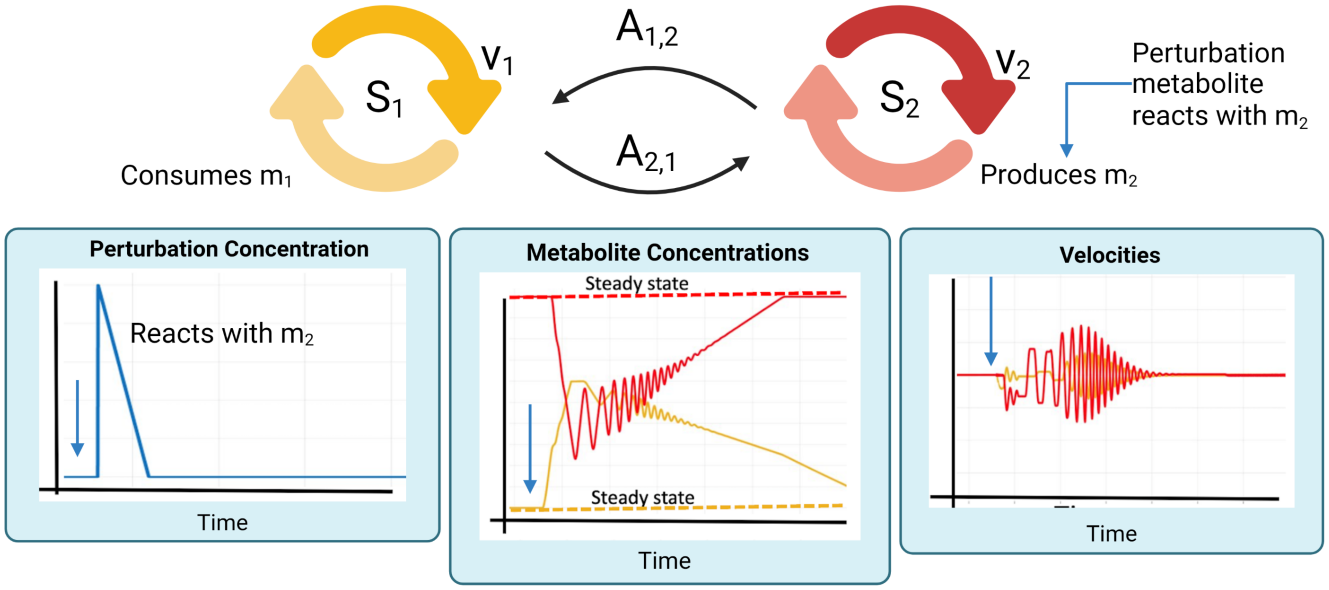
Example of two module, two metabolite system: Define two metabolic modules, *S*_1_ and *S*_2_, such that *S*_1_ consumes metabolite *m*_1_ and *S*_2_ produces metabolite *m*_2_. We can perturb the system with a toxin that is detoxified using *m*_2_. The system is then run forward in time until it converges back to its steady state.

**Fig 14.**
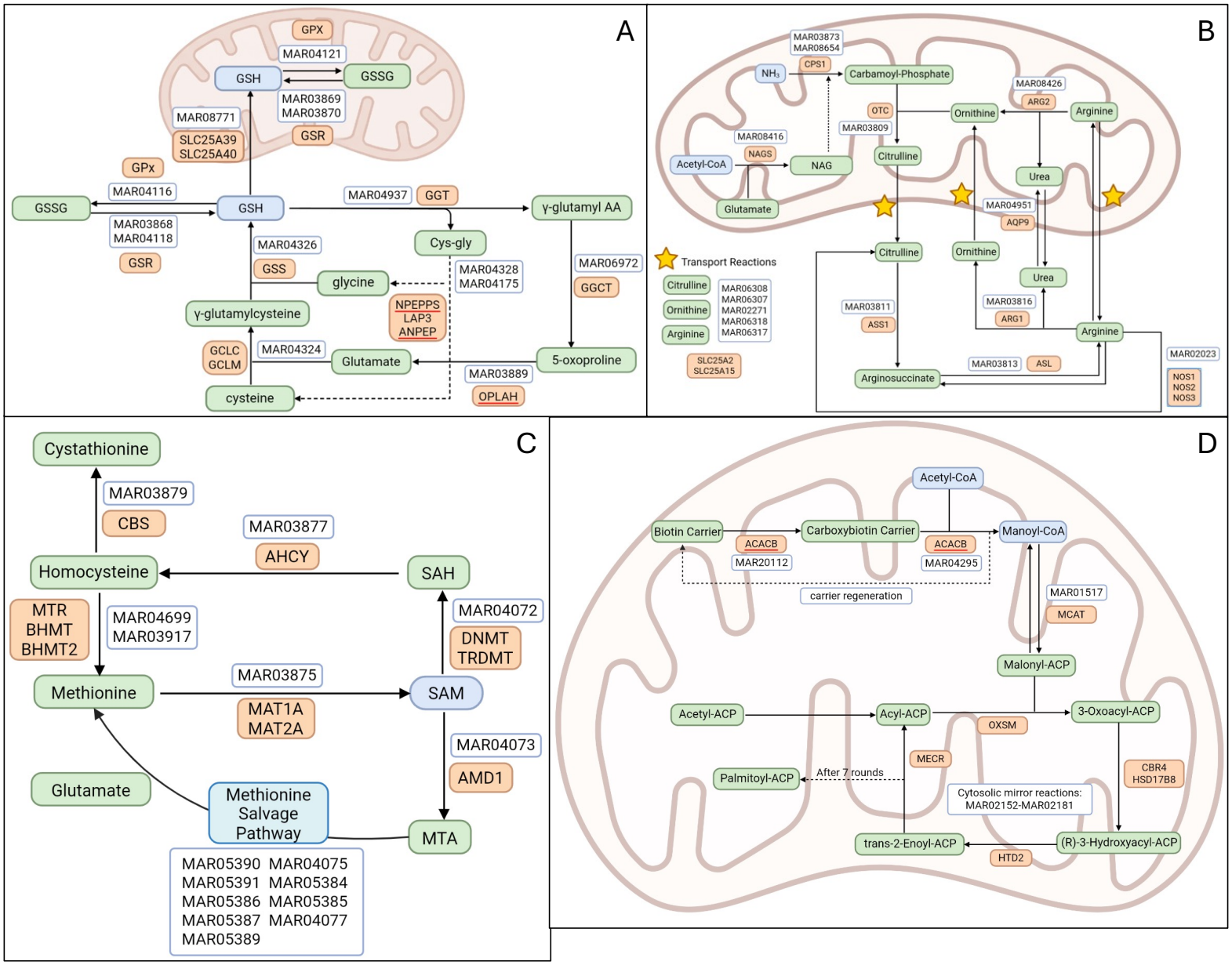
Biochemical reaction networks of each studied cycle. **(A)** The GSH cycle. **(B)** The urea cycle. **(C)** The MET cycle. **(D)** The mitochondrial fatty acid biosynthesis (FABs) cycle. Note that because the human metabolic atlas does not include the mitochondrial elongation reactions [24], we represent mitochondrial FABs using the mirrored reactants, products, and stoichiometry of the corresponding cytosolic reactions [49]. This substitution changes the catalyzing enzymes but not the reaction stoichiometry, which is the only information that enters the coupling matrix *L* and the adjacency matrix *A*.

#### 3.2.3 Critically high and low dynamics

In reality all metabolites have critically high and low concentration values. Within the context of this model, it is unnecessary to make this extension because the large separation of scales between a critically high or low value for the modeled metabolites. For example, the normal glutathione concentration in a healthy hepatocyte is approximately 10mM [46]. Glutathione is known to become hepatotoxic in a hepatocyte at concentrations that are diminished by about 70% [10] and have critical systemic impacts at about 50% depletion [47]. These concentrations are approximately 3mM and 5mM respectively. In contrast, glutathione is never found in excess as both transport and feedback loops prevent an over accumulation; note, it is even difficult to produce cells with excess glutathione in laboratory settings [48]. Similar arguments can be made for each of the metabolites we aim to model.

### 3.3 Model assumptions

The following are the assumptions taken that together define the steady state operating point around which perturbations are studied.

#### Balanced baseline

To remain viable, biochemical systems must not accumulate metabolites indefinitely. Baseline production and consumption rates are therefore assumed to balance, such that no metabolite accumulates under unperturbed conditions.

#### Comparable rates

Because metabolic subsystems are interconnected, rates across subsystems are assumed to be of comparable order of magnitude. This prevents pathological behavior in which a single subsystem dominates system wide dynamics purely by scale.

#### Fixed resource budget

The total resources of the system are assumed invariant, reflecting a fixed cellular resource budget enzyme availability, energy, redox capacity over the acute timescale considered. Consequently, our construction conserves the total velocity across all subsystems and, in the long-time limit, the total velocity associated with each metabolite.

#### Timescale separation

Fast biochemical processes are assumed to enforce approximate mass balance locally, so that the reduced system need only capture those deviations that impact flux redistribution rather than resolve detailed reaction kinetics.

#### Metabolite variables as load signals

Consistent with the flux-centric formulation, the metabolite variables [*m*_*i*_] are not interpreted as exact concentrations but as coarse-grained control signals that quantify the strain imposed on subsystems and thereby modulate resource allocation across the network. For consumed metabolites, [*m*_*j*_] represents the demand pressure exerted on consuming pathways rather than explicit depletion, reflecting upstream processes that buffer concentrations while still transmitting stress signals to the network; for produced metabolites, [*m*_*j*_] represents the accumulation pressure that drives redistribution, without requiring explicit modeling of buildup, consistent with the presence of clearance and export mechanisms.

Our approach is distinct from classical kinetic and constraint based metabolic models by shifting the primary object of interest from metabolite concentrations to perturbations of metabolic fluxes. Rather than resolving first order dynamics or the time evolution of concentrations via detailed reaction kinetics, we instead aim to capture second order effects: how the velocities of metabolic subsystems are redistributed under perturbation.

In standard kinetic formulations, metabolite concentrations *m* are treated as the principal state variables, and fluxes arise as functions *v* = *v*(*m*), yielding systems of the form 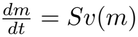, where *S* is the stoichiometric matrix. In contrast, our approach inverts this perspective.

We treat subsystem fluxes (or allocation variables) as the primary dynamical quantities and model how they are modulated by metabolite induced imbalances. The variables [*m*_*j*_] are therefore not interpreted as exact concentrations, but as effective load parameters that encode how metabolites perturb flux redistribution. Concretely, metabolite variables enter only through their influence on the resource allocation structure:

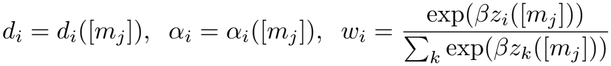

which together determine how flux is reallocated across competing subsystems. The resulting dynamics describe how perturbations propagate through the network of fluxes, rather than through explicit mass balances. From this perspective, the model captures a different level of biological organization. First order models resolve how concentrations change given fixed reaction mechanisms; here, we instead resolve how the reaction network itself reorganizes its activity in response to stress. This is particularly relevant in settings such as alcohol-associated liver disease, where pathology is characterized less by instantaneous concentration changes and more by sustained, system wide reprogramming of metabolic fluxes.

This shift justifies the interpretation of [*m*_*j*_] as coarse-grained variables:

1. For consumed metabolites, [*m*_*j*_] represents the degree of demand pressure exerted on consuming pathways, rather than explicit depletion, reflecting the fact that upstream processes act to buffer concentrations while still transmitting stress signals to the network.
2. For produced metabolites, [*m*_*j*_] represents accumulation pressure that drives redistribution, without requiring explicit modeling of buildup, consistent with the presence of clearance and export mechanisms.

Thus, the model should be understood as a reduced order, flux-centric description in which metabolite variables act as control signals governing second order responses, namely, the redistribution and coupling of metabolic velocities. This abstraction enables the model to isolate the mechanisms underlying systemic metabolic reprogramming, while avoiding the need to resolve detailed kinetic parameters that are often unidentifiable at the organ scale. Due to this nuance, we will generally be reporting and plotting the normalized load each metabolite places on the system. Denote this variable

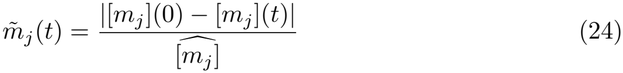

where 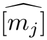 denotes the critical concentration of metabolite *m*_*j*_.

### 3.4 System specifications to model AH

Ethanol metabolism in the liver is a well studied example of how a single metabolite induces systemic metabolic perturbations through its downstream byproducts. Ethanol is oxidized via three primary pathways [35]:

1. Alcohol dehydrogenase Ethanol + NAD^+^ → Acetaldehyde + NADH + H^+^
2. Catalase Ethanol + H_2_O_2_ → Acetaldehyde + 2 H_2_O
3. Cytochrome P450 2E1 (CYP2E1) Ethanol + O_2_ + NADPH → NADP^+^ + acetaldehyde + acetate.

Acetaldehyde is a highly reactive intermediate that rapidly forms adducts with cellular macromolecules, generating products that are metabolically active or cytotoxic [35]. Its accumulation is therefore tightly controlled by the ubiquitously expressed enzyme aldehyde dehydrogenase (ALDH), which oxidizes acetaldehyde to acetate [35]:

Acetaldehyde + NAD^+^ → Acetate + NADH + H^+^.

Acetate is subsequently converted to acetyl-CoA via acetyl-CoA synthetase [35]: acetate + CoASH + ATP → acetyl – CoA + AMP + pyrophosphate.

Mitochondrial acetyl-CoA serves as a central metabolic intermediate and contributes to increased fatty acid biosynthesis, a key driver of hepatic steatosis and the development of fatty liver disease [17–19].

Under normal physiological conditions, the majority of ethanol is metabolized via alcohol dehydrogenase. The resulting excess NADH is oxidized through the mitochondrial electron transport chain, particularly at Complex I, a process that can generate reactive oxygen species (ROS) [43]. Under conditions of chronic alcohol exposure or acute high-dose intake (binge drinking), ethanol metabolism via CYP2E1 is upregulated [1, 28, 35]. Cytochrome P450 enzymes are prone to uncoupling, leading to leakage of reactive oxygen species, including H_2_O_2_ [44].

Hydrogen peroxide (H_2_O_2_) is primarily metabolized by catalase and glutathione peroxidase across cellular compartments. In the mitochondria, where our focus lies, detoxification is driven predominantly by the glutathione peroxidase pathway [42]. Maintenance of adequate mitochondrial glutathione (GSH) levels is therefore essential to prevent oxidative damage.

This highlights glutathione metabolism as a central component of the system. The glutathione pathway is tightly coupled to both the methionine cycle and the urea cycle. The methionine cycle is critical for maintaining cellular methylation capacity and regulating gene expression, processes known to be disrupted during ethanol exposure [13]. In particular the metabolite S-adenosyl-L-methionine is the principal biological methyl donor in the process [11]. In parallel, dysregulation of the urea cycle contributes to elevated blood ammonia levels, a hallmark of severe AH. Hyperammonemia is a direct driver of hepatic encephalopathy and is associated with increased morbidity and mortality in decompensated liver disease [33, 34].

Finally, increased acetyl-CoA availability promotes fatty acid synthesis, completing a mechanistic link between ethanol metabolism, oxidative stress, impaired nitrogen handling, and lipid accumulation.

In summary, ethanol metabolism is marked by an increase in ROS that is detoxed by GSH, a decrease in methylation potential, a decrease in nitrogen metabolism, and an increase in fatty acid biosynthesis. Therefore, the metabolites of interest that we will track are [SAM_*e*_]_*cyto*_, [GSH]_*mito*_, [*H*_2_*O*_2_]_*mito*_, [ammonia]_*mito*_, [acetyl-CoA]_*mito*_, and [malonyl-CoA]_*mito*_. Corresponding to our metabolites of interest, we will model the following cycles where the reaction codes are identifiable using the metabolic atlas [24].

1. Glutathione (GSH) cycle: MAR03889, MAR04324, MAR04326, MAR04328, MAR04116, MAR04121, MAR03868, MAR03869, MAR04118, MAR03870, MAR08771, MAR04937, MAR04175, and MAR06972(Fig.14(A)) .
2. Urea cycle: MAR08416, MAR03873, MAR03809, MAR02271, MAR03816, MAR04951, MAR08426, MAR03813, MAR03811, MAR02023, MAR08654, MAR06308, MAR06307, MAR06318, and MAR06317(Fig.14(B)).
3. Methionine (MET) cycle: MAR04075, MAR04077, MAR05384, MAR05385, MAR05386, MAR05387, MAR05389, MAR05390, MAR05391, MAR04073, MAR04699, MAR03875, MAR03877, MAR03917, MAR04072, and MAR03879 (Fig.14(C)).
4. Mitochondrial fatty acid biosynthesis (even chain): MAR02152, MAR02153, MAR02154, MAR02155, MAR02156, MAR02157, MAR02158, MAR02159, MAR02160, MAR02161, MAR02162, MAR02163, MAR02164, MAR02165, MAR02166, MAR02167, MAR02168, MAR02169, MAR02170, MAR02171, MAR02173, MAR02174, MAR02175, MAR02176, MAR02178, MAR02179, MAR02180, MAR02181, MAR04295, MAR01517, MAR07672, MAR07673, and MAR20112. Mitochondrial fatty acid synthesis (FABs) proceeds through the same chemistry as its cytosolic counterpart: in each round of elongation, malonyl-ACP is condensed with a growing acyl-ACP, and two NADPH-dependent reductions and a dehydration return the extended chain to a fully saturated state, adding two carbons per cycle up to chain lengths of 14-16 carbons [49]. The two pathways differ chiefly in enzyme organization. In the cytosol, each step is a catalytic domain of the multifunctional enzyme FASN, whereas mitochondrial

FABs is a type II system in which each step is carried out by a discrete enzyme (MCAT, OXSM, CBR4/HSD17B8, HTD2, and MECR) acting on the mitochondrial acyl carrier protein [49]. Because the human metabolic atlas does not include the mitochondrial elongation reactions [24], we represent mitochondrial FABs using the reactants, products, and stoichiometry of the corresponding cytosolic reactions. This substitution changes the catalyzing enzymes but not the reaction stoichiometry, which is the only information that enters the coupling matrix *L* and the adjacency matrix *A* (Fig.14(D)).

The calculation of these cycles’ corresponding adjacency matrix can be found in the supplemental section. The adjacency matrix between the above metabolic cycles is as follows:

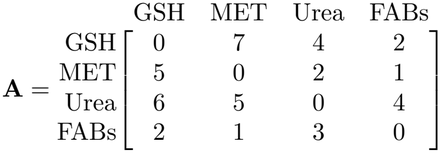

Alcohol is introduced as an exogenous, time dependent input that generates reactive oxygen species and downstream metabolic intermediates. Ethanol metabolism produces mitochondrial hydrogen peroxide via a yield coefficient that captures ethanol metabolism by CYP2E1, with this coefficient increasing under ALD conditions. Ethanol metabolism also produces downstream mitochondrial acetyl-CoA via a yield coefficient that captures altered transport of intermediate metabolites under ALD progression.

More specifically, let *S*_1_ denote the glutathione cycle subsystem, *S*_2_ denote the methionine cycle subsystem, *S*_3_ denote the urea cycle subsystem, and *S*_4_ even-chain fatty acid biosynthesis. Recall, the metabolites of interest that we will track are [SAM_*e*_]_*cyto*_, [GSH]_*mito*_, [*H*_2_*O*_2_]_*mito*_, [ammonia]_*mito*_, [acetyl-CoA]_*mito*_, and [malonyl-CoA]_*mito*_. We then have the following correspondence:

1. **Glutathione cycle**: produces [GSH]_*mito*_.
2. **Methionine cycle**: produces [SAM_*e*_]_*cyto*_.
3. **Urea cycle**: consumes [acetyl-CoA]_*mito*_ and consumes [ammonia]_*mito*_.
4. **FABs cycle**: consumes [acetyl-CoA]_*mito*_ and consumes [malonyl-CoA]_*mito*_.

Note that the detoxification of reactive oxygen species (ROS) in the mitochondria uses [GSH]_*mito*_ and is encoded in the model using third order kinetics. Now, we must define how an individual consuming an alcoholic drink perturbs the concentrations of the above metabolites. Let *a* denote the concentration of alcohol in an individual’s system. Alcohol is metabolized by the liver at a constant rate of on average .05*mM* (min^−1^), following zero order kinetics [50, 51]. Thus, we have the following differential equation for ethanol metabolism:

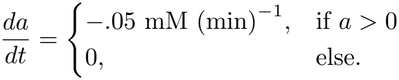

Individual genetic variation can alter this rate, as well as the other patient-specific parameters, offering a promising avenue for future work. For simplicity, we approximate ethanol influx as instantaneous. Let *y*_1_ denote the yield coefficient quantifying how many mM of [ROS]_*mito*_ is produced per 1 mM of ethanol metabolized and *y*_2_ denote the yield coefficient quantifying how mM of [acetyl-CoA]_*mito*_ is produced per 1 mM of ethanol. To capture the observed delay in mitochondrial acetyl-CoA production, we explicitly model acetate as an intermediate. Acetaldehyde, by contrast, is oxidized on a timescale much faster than the modeled processes, so we coarse-grain this effect; any ROS generated during its oxidation is absorbed into the yield coefficient *y*_1_.

Thus, we have the following differential equations, taking into account the system’s clearance dynamics:

#### Dynamic balance of [**GSH**]_*mito*_

Let *τ* denote the forward rate of the biochemical equation 2GSH + *H*_2_*O*_2_ → GSSG + *H*_2_*O*. Note that we assume the forward rate is arbitrarily large and the backward rate to be arbitrarily small given that [GSSG]_*mito*_ *<<* [GSH]_*mito*_ in general.

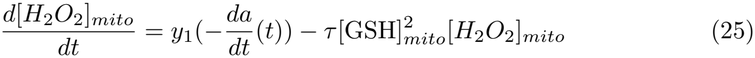

Let 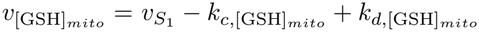.

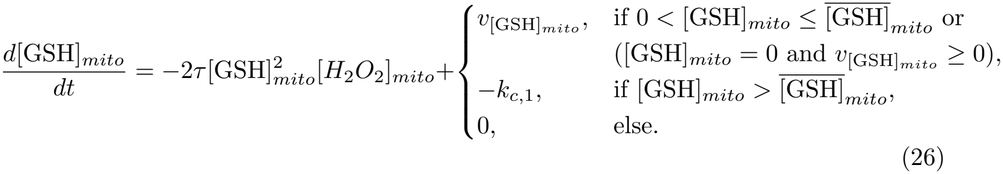

#### Dynamic balance of [**acetate**]_*mito*_

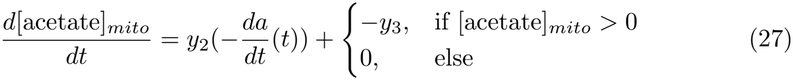

#### Dynamic balance of [**acetyl-CoA**]_*mito*_

Let 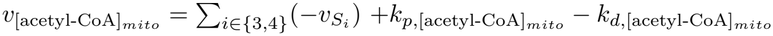.

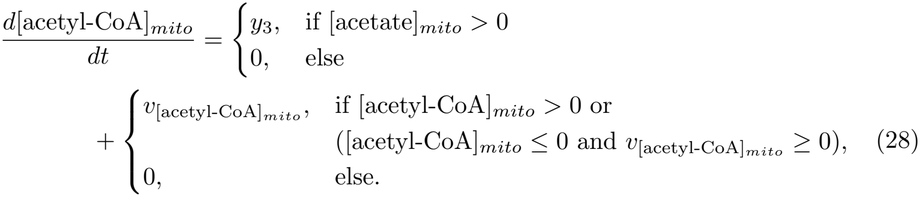

#### Dynamic balance of [**SAM**_*e*_]_*cyto*_

Let 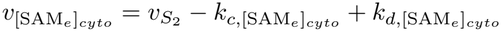

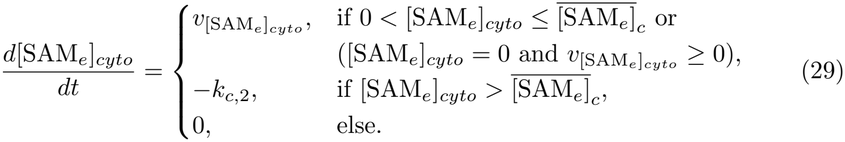

#### Dynamic balance of [**malonyl-CoA**]_*mito*_

Let 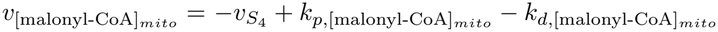

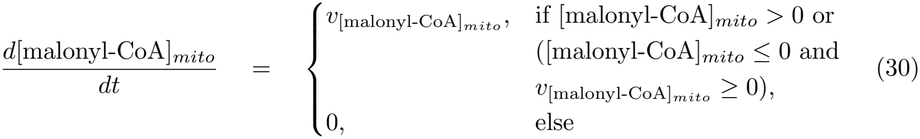

By embedding disease specific biochemical structure within a general resource allocation framework, the model provides a mechanistic bridge between ethanol exposure patterns and emergent metabolic dysfunction in AH (Fig.1).

### 3.5 Data collection, analysis, and parameter estimation

#### 3.5.1 RNA sequencing data

The transcriptomic data used in this study was previously collected and used in a previous study of ours [22]. A total of 90 human liver biopsy samples were obtained for that study as per the “Human Biorepository Core from the NIH funded international InTeam consortium” (7U01AA021908-05). The 90 patient samples were stratified into seven distinct disease states and a healthy control state: explant alcohol-associated hepatitis (explant AH, *n* = 10), severe alcohol-associated hepatitis (severe AH, *n* = 18), non-severe alcohol-associated hepatitis (non-severe AH, *n* = 11), early alcohol-associated steatohepatitis (early ASH, *n* = 12), healthy (*n* = 10), compensated cirrhosis (comp. cirrhosis, n = 9), hepatitis C virus (HCV, *n* = 10), and non-alcohol-associated steatohepatitis (NASH, *n* = 9). Patients had a median age of 45 in the explant AH disease state category, 51 in severe AH, 46 in non-severe AH, 52 in early ASH, 32 in the healthy controls, 61 in compensated cirrhosis, 46 in HCV, and 49 in NASH. Samples classified as explant AH originated from patients undergoing a liver transplant. At the time of transplant, the liver was removed from the body and was identified as having AH. A biopsy sample was then taken from the explanted liver.

Samples exhibiting severe AH were clinically and pathology proven AH and had a MELD score *>* 21. The non-severe AH patients were heterogeneous in that some displayed AH criteria (criteria discussed above) while some did not, but all had a MELD score *<* 21. The early ASH patients were non-obese, actively misused alcohol, and displayed liver damage (elevation of liver enzymes) but did not have liver dysfunction (i.e., asymptomatic, normal bilirubin, normal prothrombin levels). They also presented mild elevation of transaminases and histologic criteria of ASH but never had AH episodes. The control samples were taken before any clamping or ischemia from the normal liver surrounding apparent liver nodules as a result of metastases. The patient samples from the compensated cirrhosis, HCV, and NASH cohorts were all clinically and pathology proven proven, and none of the patients had any AH episodes. Patients from the compensated cirrhosis cohort were asymptomatic and had HCV and cirrhosis but did not have ascites, variceal hemorrhage, hepatic encephalopathy, or jaundice. Patients within the HCV cohort did not have cirrhosis, and patients within the NASH cohort were defined according to Keiner’s Criteria and displayed accumulated hepatic fat histologically and had above average body mass index (BMI *>* 25). Patient inclusion and exclusion criteria in the clinical trial, as well as additional information about the samples, can be found on the clinical trial website (https://clinicaltrials.gov/study/NCT02075918).

#### 3.5.2 Fluxome prediction

Patient specific metabolic fluxomes were predicted using the Pheflux algorithm [52] as used in our previous work [23]. This methodology seeks to infer the most likely and least biased fluxome by maximizing Shannon’s entropy of fluxes per mRNA without the need for an objective function. Maximization of Shannon’s entropy is equivalent to minimizing the forward Kullback-Leiber divergence between the fluxome and transcriptome. The input to the Pheflux algorithm is the CPM normalized RNAseq data and a reference metabolic network, for which we used the Human1 network [24]. The output is the patient specific metabolic fluxomes, one flux for each reaction in the Human1 network. The Pheflux algorithm was implemented in Python using the source code found in the author’s GitHub repository: https://github.com/mrivas/pheflux. Nonzero fluxes were identified as those greater than zero for all patients for a given reaction. An empirical cumulative distribution function (ecdf) plot was generated for the nonzero log10 transformed metabolic fluxes, and the knee point was established as the cutoff for reactions with significant flux values. The coordinates of the knee point are (0.001, 0.534), corresponding to a flux cutoff at 0.001 mmol/gDW/g, which consists of 53.4% of the entire fluxome. The ecdf function from the stats package in RStudio was utilized to calculate the ecdf, and ggplot2 was utilized for plotting.

### 3.6 Parameter estimation and numerical implementation

#### Initial velocities per cycle

This modeling framework conserves system resources and therefore, by construction, preserves the total velocity. Without loss of generality, we set the total velocity 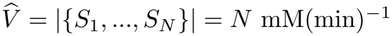. To compute the module specific proportions of the total velocity we start with use RNA sequencing data from individuals under various stages of ALD progression as described in Sec. 3.5.1. We use this data in genome scale metabolic modeling to predict individual fluxomes following our published approach in [52] summarized in Sec. 3.5.2. We then found the average flux through each metabolic module over each disease state. Denote these averages fluxes as 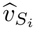 for metabolic module *S*_*i*_. We compute the normalized initial metabolic velocity as 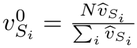.

The fluxome predictions yielded the initial velocities across four different disease states:

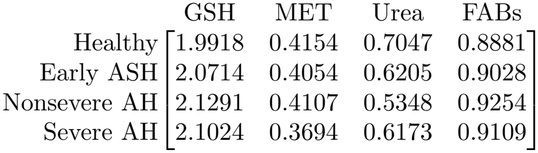

Since these initial velocity estimates were similar across disease states, in the subsequent analysis, we use an average as

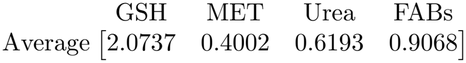

where the units are mM/(min).

#### Initial concentrations

For the metabolites that are consumed by their associated metabolic module, the initial conditions were shifted to a concentration 0 mM. In contrast for metabolites that are produced, the initial concentrations were set to available experimental measurements. Specifically, we use .08 mM as the initial concentration of cytostolic SAMe [11, 53] and 10 mM as the initial concentration of mitochondrial glutathione [46].

#### Critical concentrations 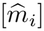

The critical concentration of the modeled metabolites were deciphered based on available literature:

1. 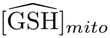 A 50% depletion of mitochondrial glutathione is considered as a pathological state [47]. Therefore, we consider the mitochondrial GSH critical value to be halfway between the pathological level and its initial concentration, ie 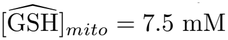.
2. 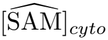 A 40% − 60% decrease in 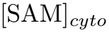 causes a noticeable shift but not pathological in methlyation capacity [54]. Therefore, we consider 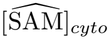 to be at 50% the initial concentration, ie 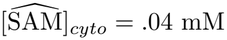 mM.
3. 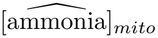 Ammonia concentrations above 5 mM cause collapse of mitochondrial membrane [55]. Additionally, alterations of mitochondrial dehydrogenase have been shown to start becoming noticeable at approximately 1 mM ammonia [55]. Therefore, we will take the critical concentration of ammonia to be 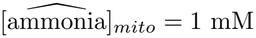.
4. 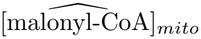 Precise hepatotoxic concentrations of mitochondrial malonyl-CoA are not readily established, as a wide range of values has been suggested [56]. One landmark study achieved approximately 25% inhibition of fatty acid oxidation using roughly 30 *µM* mitochondrial malonyl-CoA in starved rats [56, 57]. To approximate the midpoint of progression toward pathology, we instead seek the concentration producing 50% inhibition. Given the shape of the dose response curve in [57], we estimate this concentration to be roughly 10*×* the amount used above, or 300 *µM* = 0.3 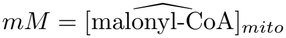. We note that we adopt the starved rat concentrations from [57] because ALD and ethanol consumption are known to be associated with malnutrition [58].
5. 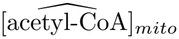 Like mitochondrial malonyl-CoA, the hepatotoxic concentrations of mitochondrial acetyl-CoA is not known in literature. Unlike mitochondrial malonyl-CoA, there do not exist estimates of this number. Because mitochondrial acetyl-CoA is a common biologcial intermediate molecule [59], we take it to be double the highest of the consumption based metabolites, [ammonia]_*mito*_. Thus, 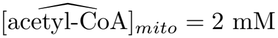.

#### Time to detox (Δ*t*)

The time to detox was chosen to be approximately 1 hour which is the amount of time it takes an average individual to detox approximately one US standard drink of alcohol [50].

#### Yield coefficients *y*_1_, *y*_2_, *y*_3_

The parameter *y*_3_ is taken to be 1 mM as the time scale of the reaction it models is relatively short and should not vary across disease state. In contrast, the yield coefficients *y*_1_ and *y*_2_ are patient specific parameters that are allowed to vary across disease state. Recall that *y*_1_ is the yield coefficient converting the mM ethanol metabolized to the mM mitochondrial H_2_O_2_ produced. Mitochondrial H_2_O_2_ is produced at various stages of ethanol metabolism and subsequent downstream processes including ethanol oxidation by CYP2E1 and acetaldehyde oxidation.

Immediately following ethanol consumption there is an extreme depletion of GSH [30] with a time scale of a few hours. Note that this time scale is the same as that of ethanol oxidation to acetaldehyde. Given 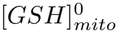 is on the order of 1 mM and 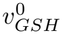 is on the order of 1 mM(min)^−1^ (see initial velocity calculation in this section), *y*_1_ should be between the scales of .01 mM to 1 mM. Thus, we will take *y*_1_ ∈ [.01, .75] mM. Similarly, *y*_2_ is the amount of downstream metabolites that are retained in the mitochondria that eventually become mitochondrial acetyl-CoA. According to the RNA-seq data and corresponding predicted fluxomes described in Sec. 3.5, the transport of acetate out of the mitochondria under ALD progression is significantly down regulated (SI.3.7). This implies that *y*_2_ should increase under ALD progression. Further *y*_2_ should always be bound by 1mM as more acetyl-CoA cannot be produced than ethanol consumed given the biochemical network reactions. Thus, we take *y*_2_ ∈ [.1, 1] mM.

#### Time step

In order to solve these equations, we implemented Forward Euler in time. We used a time step *dt* = .005 minutes to maintain stability and eliminate spurious oscillations that occur as a result of larger time step values. Note that because *τ >>* 1, we put a stop gap so that the simulation never enters a negative concentration regime without having to reduce *dt* substantially in order to resolve these fast time scale dynamics.

#### Other values

Additional values are taken to be as follows:

**Table 1.**
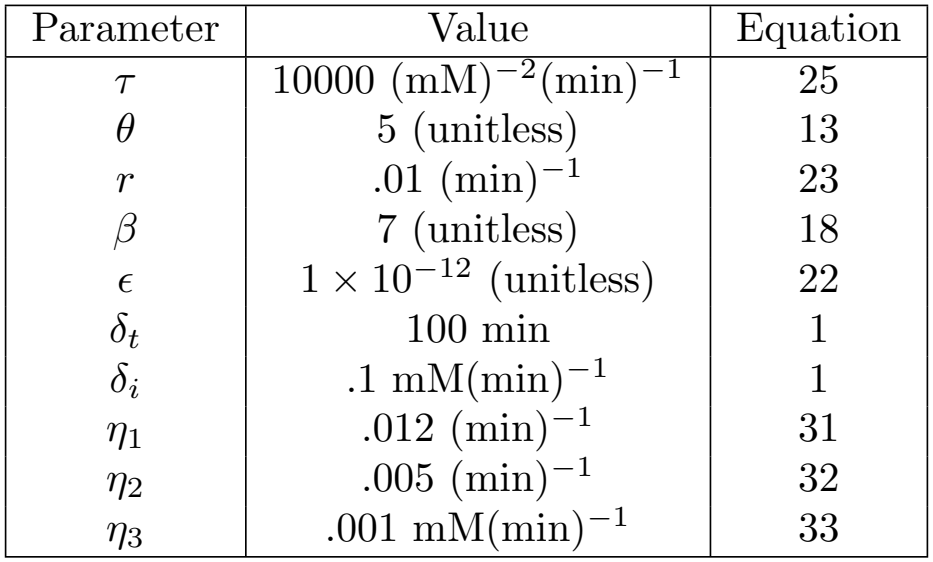
Table of addtional parameter: Table of additional noted parameter values not reported above.

| Parameter | Value | Equation |
|---|---|---|
| *τ* | 10000 (mM)<sup>−2</sup>(min)<sup>−1</sup> | 25 |
| *θ* | 5 (unitless) | 13 |
| *r* | .01 (min)<sup>−1</sup> | 23 |
| *β* | 7 (unitless) | 18 |
| *ϵ* | 1 × 10<sup>−12</sup> (unitless) | 22 |
| *δ<sub>t</sub>* | 100 min | 1 |
| *δ<sub>i</sub>* | .1 mM(min)<sup>−1</sup> | 1 |
| *η*<sub>1</sub> | .012 (min)<sup>−1</sup> | 31 |
| *η*<sub>2</sub> | .005 (min)<sup>−1</sup> | 32 |
| *η*<sub>3</sub> | .001 mM(min)<sup>−1</sup> | 33 |

### 3.7 Numerical Experimental Procedures

Except where noted, results derive from a common simulation ensemble. We vary the three patient specific coefficients: the ROS yield *y*_1_, the acetyl-CoA yield *y*_2_, and the transport coefficient *k*_*d*_ over their physiological ranges (Sec. 3.6) on a 11 *×* 11 *×* 11 grid, with *y*_1_ ∈ [.01, .75], *y*_2_ ∈ [.1, 1], and *k*_*d*_ ∈ [.001, .02] varying uniformly. For simplicity of analysis we set 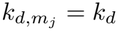 equal for all metabolites.

#### Canonical phenotypes scenarios

To assess clinically observed attributes, we define two numerical experiments, each simulated over 4520 minutes post ethanol exposure (approximately 72 hours). In the one-hit experiment, an ethanol perturbation of 17.4 mM is applied at *t* = 200, after which the system evolves freely toward its perturbed steady state. In the two-hit experiment, the same perturbation is applied at *t* = 200 and again at *t* = 4520 minutes (72 hours later). For each experiment we compute the net deviation of module velocities from their steady state values over the full window, measuring how resources are redistributed across cycles, and we track 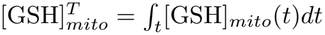 and [H_2_O_2_]_*mito*_. Each simulation is run by sweeping the previously listed patient specific values over a 11 *×* 11 *×* 11 grid.

#### Acute GSH depletion in mouse model experiment

We adjust the model to mouse metabolism, changing the ethanol clearance rate from .05 to .275 mM(min)^−1^, reflecting the roughly 5.5*×* higher capacity of mice [60]. Following the experiment conducted by [30], we administer a 5 g/kg dose which corresponds to approximately 100 mM given a 0.02-0.035 kg animal. For a healthy, fully matured 12 month old mouse we take *y*_1_ = 0.06, *y*_2_ = 0.15, *k*_*d*_ = 0.01. The dynamics are allowed to rest at equilibrium until 100 mM ethanol at *t* = 200 min, and integrate forward for 24 hour simulation time (to *t* = 1640 min). We still retain the initial condition of [GSH]_*mito*_ = 10 mM.

#### Heterogeneous failure scenario

We examine representative healthy, diseased (ALD), failed-compensated (AH), and failed-decompensated (AH) individuals, each receiving a 17.4 mM perturbation at *t* = 200 over a 3200 minute simulation. Parameters: healthy (*k*_*d*_, *y*_1_, *y*_2_) = (.01, .2, .25); diseased (.005, .5, .6); failed-compensated (.003, .75, .5); failed-decompensated (.001, .75, .6).

#### Cumulative exposure sensitivity analysis

Variance based sensitivity analysis on 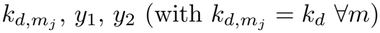 was preformed by estimating first order and total effect indices via [32, 61] at *N* = 1024 = 2^10^. Two scenarios are compared: a one-hit (17.4 mM) and a two-hit (17.4 mM twice, separated by 240 min).

#### Corticosteroid treatment scenario

We represent corticosteroid treatment as a reduction in the effective glutathione consumption rate 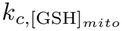 not a direct elimination of ROS, but a reduction in inflammatory/oxidative burden and thus in the effective demand on mitochondrial GSH buffering. Treatment acts as a temporary crutch, allowing resources to flow to other metabolic modules that would otherwise have been taken by the GSH cycle. We take the 72 hour ensemble and identify AH trajectories as those in *M*_*a*_ persistently metabolically decompensated at terminal time.

All treatment simulations begin with a single 17.4 mM perturbation at *t* = 200 min, with the horizon extended to *T* = 25,000 min *≈* 17 days. Treatment is modeled phenomenologically via an internal corticosteroid burden *S*(*t*) and cumulative effect *E*(*t*):

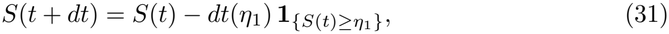

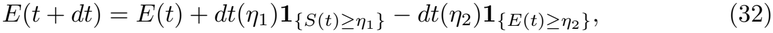

so active exposure generates a transient effect of corticosteroids that decays on a slower timescale. Both *η*_1_ and *η*_2_ are tuned so the effect enters within the first hour and leaves within 36 hours of administration, consistent with known corticosteroid pharmacokinetics [62, 63]. The effect of corticosteroids couples into the GSH subsystem by reducing baseline consumption:

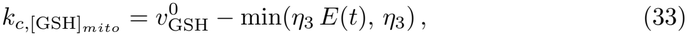

with 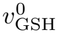 the baseline glutathione velocity and *η*_3_ chosen to match the smallest admissible *k*_*d*_, keeping treatment effects physiologically constrained.

Three intervention schedules were considered, each with seven daily administrations (1440 min interval): early (*t* = 2880 min *≈* 48 h), intermediate (*t* = 4320 min *≈* 72 h), and late (*t* = 5760 min *≈* 96 h). Efficacy was evaluated at day 17: realizations were classified metabolically unstable if the final deviation exceeded the threshold criterion, and schedules were compared by their ability to restore stability in otherwise decompensated trajectories. We quantify long time instability of a trajectory *x*(*t*) through

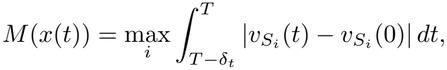

the cumulative terminal window deviation of subsystem velocities from baseline. For a treatment regimen *φ*, we say treatment produces a net stabilizing effect if *M* (*φ*(*x*(*t*))) *< M* (*x*(*t*)).

#### Threshold failure scenarios

We vary the three patient specific coefficients: the ROS yield *y*_1_, the acetyl-CoA yield *y*_2_, and the transport coefficient *k*_*d*_ over their physiological ranges (Sec. 3.6) on a 11 *×* 11 *×* 11 grid, with *y*_1_ ∈ [.01, .75], *y*_2_ ∈ [.25, 1], and *k*_*d*_ ∈ [.001, .02] varying uniformly. We probe the parameter space under multiple perturbations across four scenarios:

1. **Single hit:** 17.4 mM ethanol added.
2. **Double hit:** 17.4 mM added, then a further 17.4 mM after 60 min.
3. **Triple hit:** as double hit, then a further 17.4 mM after 24 h (1440 min).
4. **Quadruple hit:** as triple hit, then a further 17.4 mM after another 60 min. (Fig. 10)

Results are labeled as pass or failed for each of the four scenarios where failure indicates a nonzero transient spike of [H_2_O_2_].

## Acknowledgments

The authors would like to acknowledge financial support for this study from the National Institute on Alcohol Abuse and Alcoholism R01 AA018873 (PIs: R.V.). The funding sponsors had no role in the design of the study; in the collection, analyses, or interpretation of data; in the writing of the manuscript, and in the decision to publish the results.

## Data availability and reproducibility

All data and associated code for the model simulations and analysis are available in the main text or the supporting information. There are additionally hosted on Github at: https://github.com/Madison-Shoraka/AH-Metabolic-Modeling. To test the reproducibility, Claude Opus 5.5 (Anthropic) was provided with the manuscript PDF, in a session with no access to the original source code, and asked to independently implement the model. The resulting code correctly reproduced the disease states shown in Fig. 4. These results were obtained with *reasoning-effort* parameter was HIGH, as lower settings were unable to reproduce the results. The Claude-generated code is included in the supplemental information for comparison (SI. 3.7). In the above GitHub repository, we provide a self-assessment of credible practice for modeling and simulation in health care following the ten simple rules specified in [64] with the evaluation rubric found in [65].

## AI use statement

To test the reproducibility, Claude Opus 5.5 (Anthropic) was provided with the manuscript PDF, in a session with no access to the original source code, and asked to independently implement the model. The resulting code correctly reproduced the disease states shown in Fig. 4. That implementation is available in the GitHub repository listed under Data Availability for comparison. The mathematical formalism and all scientific ideas presented here are the authors’ own. Claude was not used to generate the original model code, but was used to optimize run time and matrix implementation and to assist with debugging. The authors take full responsibility for the content of this manuscript.

## Supporting information

- **S1 Code. Code and results file associated with Sec. 2.1, 2.3, 2.4:** This folder contains code and resulting data files for original grid search in Sec. 2.1, entire Sec. 2.3, and percentage of cases in failure in Sec. 2.4.
- **S2 Code. Code and results file for network analysis in Sec. 2.1 and simulation in Sec. 2.3:** This folder contains code and resulting data files for network grid search in Sec. 2.1 as well as the example simulations found in Sec. 2.3.
- **S3 Code. Code and results file for UQ found in Sec. 2.4:** This folder contains code and resulting data files for uncertainty quantification found in Sec. 2.4.
- **S4 Code. Code and results file for simulations found in Sec. 2.5:** This folder contains code and resulting data files for the treatment analysis found in Sec. 2.5.
- **S5 Code. Code and results file for simulations found in Sec. 2.6:** This folder contains code and resulting data files for the multi-hit stability analysis found in Sec. 2.6.
- **S1 Data. Data and resulting analysis used for patient specific parameters:** This folder contains data and resulting analysis used for the computation of the stoichiometry coupling and initial velocities as seen in Sec. 3.6.
- **S6 Code. Code mentioned in Reproducibility section** This file contains code mentioned in the reproducibility section.

